# Establishment of cell transcriptional identity during seed germination

**DOI:** 10.1101/2023.01.21.523180

**Authors:** Lim Chee Liew, Yue You, Marina Oliva, Marta Peirats-Llobet, Sophia Ng, Muluneh Tamiru-Oli, Oliver Berkowitz, Uyen Vu Thuy Hong, Asha Haslem, Tim Stuart, Matthew E. Ritchie, George W. Bassel, Ryan Lister, James Whelan, Quentin Gouil, Mathew G. Lewsey

## Abstract

Germination involves highly dynamic transcriptional programs as the cells of seeds reactivate and express the functions necessary to establish in the environment. Individual cell types have distinct roles within the embryo, so must therefore have cell-type specific gene expression and gene regulatory networks. We can better understand how the functions of different cell types are established and contribute to the embryo by determining how cell-type specific transcription begins and changes through germination. Here we describe a temporal analysis of the germinating Arabidopsis embryo at single-cell resolution. We define the highly dynamic cell-type specific patterns of gene expression and how these relate to changing cellular function as germination progresses. Underlying these are unique gene regulatory networks and transcription factor activity. We unexpectedly discover that most embryo cells transition through the same initial transcriptional state early in germination, after which cell-type specific gene expression is established. Furthermore, our analyses support previous findings that the earliest events leading to the induction of embryo growth take place in the vasculature. Overall, our study constitutes a general framework to characterise Arabidopsis cell states through embryo growth, allowing investigation of different genotypes and other plant species whose seed strategies may differ.

## Introduction

Germination is the process through which seeds begin to grow and establish in their environment, which is fundamental to agricultural production. Success requires that the seed monitor both its internal resources and surrounding, external conditions, then co-ordinate appropriate responses based upon this information to ensure that the time and place are suitable for future plant growth. Germination and early seedling growth are consequently plastic, dynamic processes. The seed is a complex structure comprised of many tissues and cell types [1–3]. These have individual functions and biochemistry that enable tight spatiotemporal control of growth and development. Variation in temperature is sensed and interpreted as a signal to germinate by just tens of cells in the embryonic radicle of dormant Arabidopsis seeds, in conjunction with the monolayer of cells in the endosperm [4, 5]. Growth is then driven by cell expansion once germination commences, with both spatial and temporal variation in expansion rates. Initially, growth occurs in cells adjacent to the radicle tip, then proceeds to include cells further along the radicle and in the hypocotyl [6]. The end of germination is defined by the emergence of the radicle through the testa (seed coat), following which the cotyledons emerge and the seedling transitions from heterotrophic to photoautotrophic growth [7].

Spatiotemporal control of growth and development requires correspondingly precise regulation of gene expression. The nuclei of mature Arabidopsis embryo cells condense and transition to a heterochromatic state by the end of seed development, repressing gene expression [8–10]. This is thought to be an adaptation to tolerate the desiccation of seeds that occurs at the end of seed development [9]. The nuclei then reverse this process as the seed imbibes water and germination commences, decondensing and transitioning to the euchromatic state required for gene transcription [9, 11]. However, germination considered in the strict sense (i.e. to the point of radicle protrusion) is thought to be dependent only on translation, whereas *de novo* transcription during germination is non-essential [12]. Consistent with this, mature Arabidopsis seeds contain populations of stored transcripts that were transcribed during seed development, a subset of which are translated early in germination [12–14]. During this time the developing embryo draws upon stored energy reserves, primarily in the form of lipids but also from cell wall carbohydrates [8, 15–18]. Nonetheless, transcription and *de novo* gene expression occur relatively early in germination, within 1-2 hours of imbibition [19, 20]. A cascade of transcription factors regulate gene expression at this time, which influences the speed at which germination progresses and is essential for a successful transition to post-germination seedling establishment [5, 13, 19, 21, 22]. Temporal changes occur in DNA methylation and small RNA abundance during germination, both of which are likely to be involved in gene regulation [13, 23, 24]. Gene expression also varies spatially within seeds, reflecting the different functions of tissues and cell types [1, 4, 25–27]. However, the resolution of spatial studies, and consequently the insight they provide, have been limited by the precision and scale achieved by the predominant methods of hand isolation or laser capture microdissection [1, 25, 26].

By understanding the spatiotemporal regulation of gene expression in seeds we can better understand how the functions of different cell types are specified, and how these contribute to the functions of the seed as a whole. To this end, we investigated gene expression dynamics in the Arabidopsis embryo over the first 48 hours of germination, as it transitions into a seedling, at single-cell resolution. We then interrogated the data to understand how the transcriptional identity of cells was established and how gene expression was regulated. We observed that most cells pass through a shared early transcriptional state, before transitioning to their cell-specific transcriptional states. Once established, these cell-specific transcriptional states were dynamic over germination and reflected changing functional properties of cells. We constructed gene regulatory models for each cell type, from which we predicted key transcription factors active in individual cell types. This study provides unprecedented insight into the different regulatory mechanisms operating as a seedling establishes itself within the environment.

## Results

### Generating a single-cell gene expression atlas of germinating embryos

The major goals of our study were to characterise how gene expression differs between the cell types that constitute germinating Arabidopsis embryos and to determine how these patterns of gene expression may be regulated. To this end, we generated a single-cell RNA-seq (scRNA-seq) atlas of germinating Arabidopsis embryos (Fig. 1). Three time points post-stratification were selected for analysis (12, 24 and 48 hours), corresponding to early, mid and the end of germination in our conditions (Fig. 1a) [13]. Seeds were harvested at each time point and embryos were released from the seed coat by physical disruption. Protoplasts were then isolated and enriched to high purity in order to physically separate the cells from one-another and make them amenable to microfluidic handling.

**Fig. 1.**
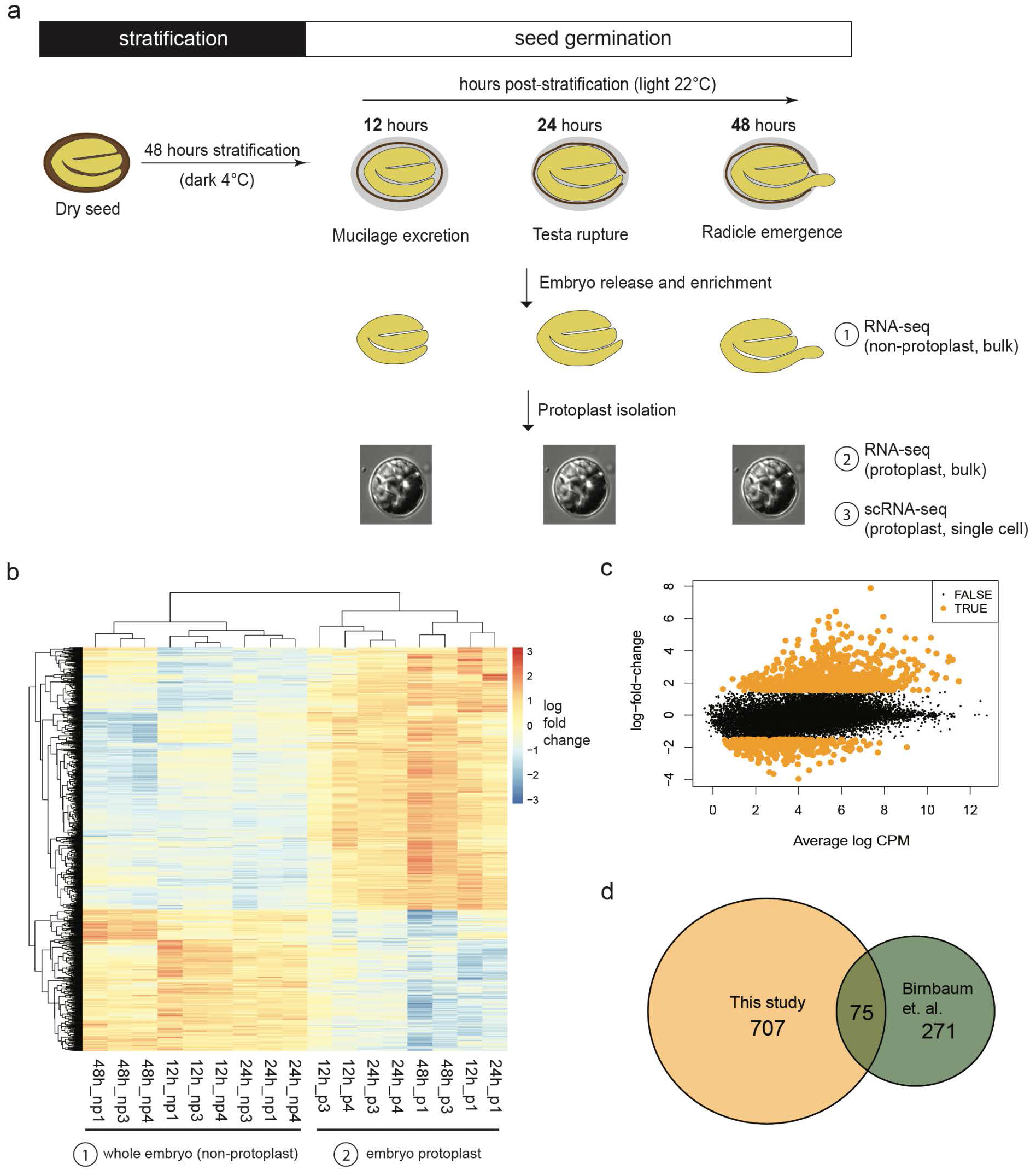
Germinating embryo scRNA-seq experimental design and impact of protoplast isolation on the transcriptome. **a,** Germination procedure and sampling for transcriptomic analyses. (1) RNA-seq of whole isolated embryo without protoplast isolation; (2) RNA-seq of embryo protoplasts; (3) scRNA-seq of individual embryo protoplasts. **b,c** Expression changes of 1,202 genes in response to protoplast isolation (log-fold changes). The transcriptional response to protoplast isolation was consistent across time-points. **d,** Limited overlap in upregulated genes upon protoplast isolation between this study and a previous study by Birnbaum and colleagues [28].

Isolation of protoplasts affects gene expression in sampled cells, creating a technical effect that must be controlled for in subsequent analyses [28]. To enable this correction we analysed the effect of protoplast isolation on embryo transcriptomes using bulk RNA-seq (i.e. not scRNA-seq). We compared transcriptomes in whole, isolated embryos with those of our protoplast preparation, finding that 1,202 genes were differentially expressed between them (1% FDR, log2 fold-change greater than 1.5, Fig. 1b-c, Supplementary Table 1). The genes responsive to protoplast isolation responded consistently across all time points (Fig. 1b). These genes were excluded from our subsequent scRNA-seq data analyses.

Other plant single-cell gene expression studies have also used protoplast isolation to separate cells [28–34]. Approaches to correct for the effect of protoplast isolation have varied; some studies have applied bulk RNA-seq in the same manner as ourselves, whilst others have either used a single previously-generated reference set of protoplast-isolation responsive genes [28] or reported no correction at all. It is possible that the population of protoplast isolation responsive genes varies dependent upon experimental conditions, growth stage and other factors. We tested this by determining how many of the responsive genes were shared between our analysis and the previously generated reference set. Only 75 of our 782 upregulated protoplast isolation responsive genes were shared between the two datasets (Fig. 1d, Supplementary Table 1). This indicates that it is important the effects of cell or protoplast isolation are analysed and controlled for in parallel when conducting scRNA-seq.

Transcriptomes of germinating embryo protoplasts were analysed using the 10x Genomics Chromium platform in order to produce a single-cell atlas (Fig. 2, Extended Data Fig. 1). Two biologically-independent replicates were conducted per time point. This yielded a total of 12,798 cells, with an average 1,025 expressed genes per cell (Extended Data Fig. 1a,b). Cell recovery and the number of detected genes increased with time of germination, though equal numbers of cells were loaded per sample (Extended Data Fig. 1a). It is likely this occurred because the amount of transcripts per cell was low in the early stages of germination, affecting the detection of true cells from background [13]. We integrated data from all samples to minimize technical effects, then clustered cells according to their transcriptional profiles, and visualised the resulting clusters in two dimensions with UMAP (uniform manifold projection) to assess consistency between replicates and time points (Fig. 2a, Extended Data Fig. 1c-e). Cells from independent replicates of individual time points were co-located in the analysis, indicating replicates were consistent with one another after data integration (Extended Data Fig. 1d). Fifteen distinct clusters of cells with similar transcriptional profiles were resolved amongst these data, which likely represent different cell types and states (Fig. 2a).

**Fig. 2.**
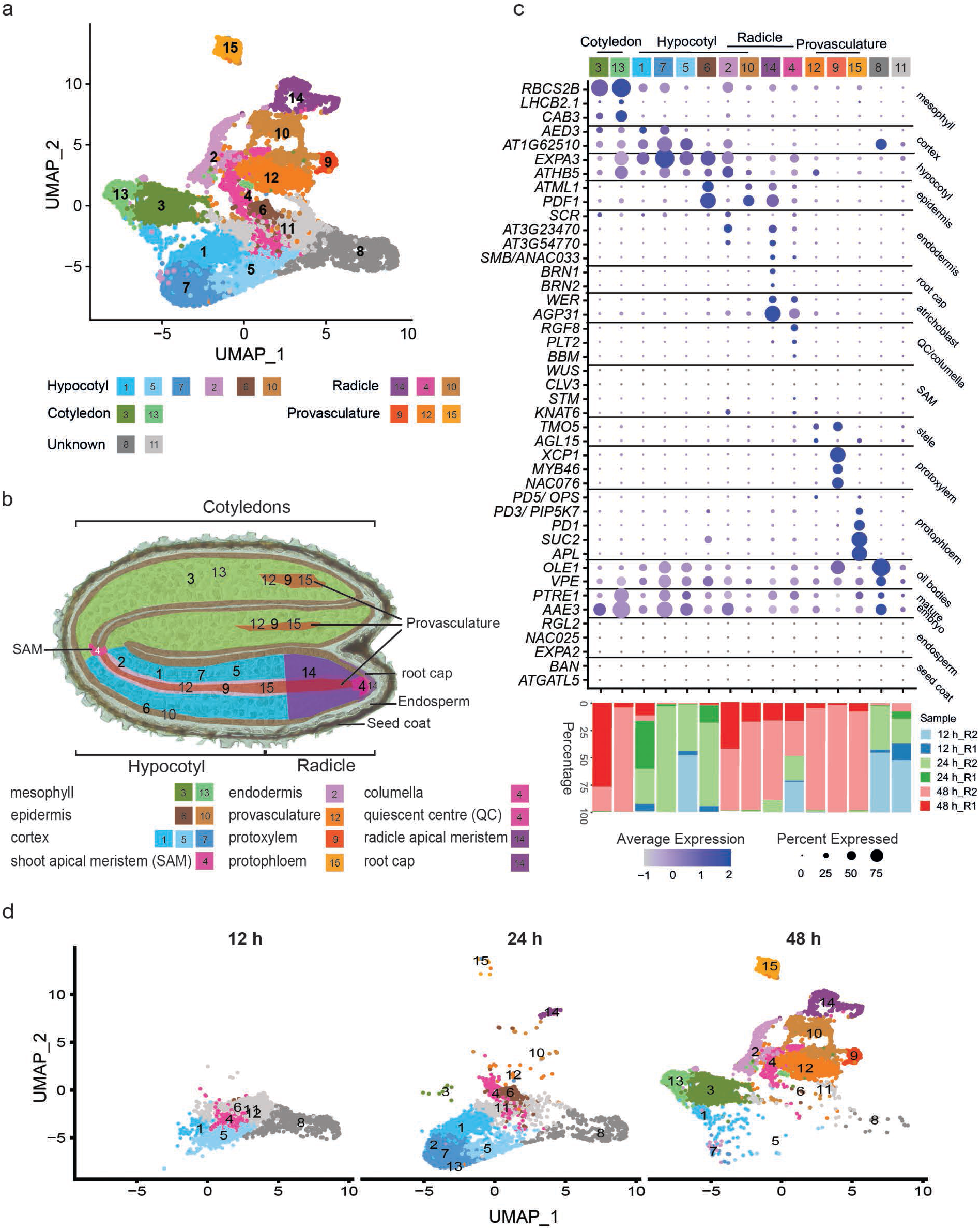
Annotation of germinating embryo cell types using literature curated marker transcripts. **a,** UMAP dimensional reduction and visualisation of 12,798 cells in 15 clusters. **b,** Spatial distribution of cell clusters in the Arabidopsis embryo. **c,** Bubble plots showing enrichment of expression of representative cell-type specific marker transcripts in 15 clusters and the percentage of cells from the two biological replicates within each cluster at each time point. Marker transcripts were identified from published studies **d,** UMAP dimensional reduction and visualisation of cells across three time points, 12 h, 24 h and 48 h, showing the temporal changes in cell and cluster detection.

The cell clusters detected within embryos differed markedly between the 12, 24 and 48 h time points (Fig. 2c, d, Extended Data Fig. 1f). No cell division occurs in embryos during germination, so no new cells arise in the time-period studied [6]. Consequently, the differences in cell clusters between time-points indicate that the transcriptional states of individual cell types within the embryo change over time during germination, even when considering that we did not capture a subset of transcript-poor cells at the early timepoints. This means that the cell clusters defined in our experiment correspond to ‘cell states’ adopted by the various cells and tissues through the timecourse of germination, rather than to cell types directly. The consistency between replicates at individual time points was considered (Fig. 1d,f). Largely the same clusters were present in each replicate, but the proportions of cells in those clusters varied. As gene expression is highly dynamic at this time, even a very small shift in harvest time or environmental conditions might cause differences between replicates. Overall, however, the replicates corresponded well with one another.

### Annotation of cell types in the germinating embryo single-cell gene expression atlas

We annotated the clusters within the embryo single-cell atlas so that we could interpret the changes occurring amongst them during germination (Fig. 2c). Annotation of single-cell data is often achieved by reference to an existing ground-truth dataset of manually dissected or sorted cells from the organ of interest [28, 30–32]. A comprehensive ground-truth dataset does not exist that describes all cell types of the embryo during germination. To overcome this we investigated the published literature and identified relevant marker genes from several studies, then compared them to genes expressed specifically in one or a small number of clusters (Supplementary Table 2).

We were able to infer identities for thirteen of the fifteen clusters (Fig. 2c). We detected the most abundant cell types of the cotyledon (mesophyll, clusters 3, 13), hypocotyl (cortex, clusters 1, 5, 7; epidermis, 6; cortex/endodermis, 2) and radicle (epidermis, 10, radicle apical meristem, 14; quiescent centre/columella, 4). Cell-type markers were expressed clearly by cluster 2 (cortex/endodermis) and cluster 10 (epidermis), but we could not distinguish whether these were resident in the hypocotyl or radicle, likely reflecting that these cell types are continuous between the two organs at this developmental stage. Cells of the provasculature (protophloem, 15; protoxylem, 9; provascular cells, 12) were mostly detected at the 48 h time point. Identities could not be assigned to clusters 8 and 11 because they did not show clear enrichment of expression for any marker genes from the literature, indicating the clusters may represent some uncharacterised cell state or type. Clusters strongly expressing the marker transcripts of the shoot apical meristem *WUSCHEL* (*WUS*, *AT2G179500*), *CLAVATA3* (*CLV3*, *AT2G27250*), *SHOOT MERISTEMLESS* (*STM*, *AT1G62360*) and *KNOTTED1-LIKE HOMEOBOX GENE 6* (*KNAT6*, *AT1G23380*) were not detectable in the dataset. Some enrichment of KNAT6 and STM was detected in clusters 4 and 10, suggesting that these clusters might include the small number of shoot apical meristem cells. The absence of a clear shoot apical meristem cluster likely occurred because these cells are very rare (approximately 8 cells) relative to the total number of cells in the embryo.

We first validated annotations for clusters 9 and 14 by RNA *in situ* hybridisation. We annotated cluster 9 as protoxylem within the provasculature, and present only at 48 h. The scRNA-seq analysis indicated that *AT1G55210* expression was an independent marker for cluster 9, which had not been used in the initial literature-based annotation of the cluster, so we determined the location of its transcripts by RNA *in situ* hybridisation (Fig. 3a, Supplementary Tables 3, 4, Extended Data Fig.2). Correspondingly, a signal was detected specifically within the protoxylem at 48 h, but was not detected at 24 h. We also validated the annotation of cluster 14 as radicle apical meristem cells, present only at 48 h. The location of the independent marker transcript *AT3G20470* was examined. A signal was specifically detected in cells of the radicle apical meristem region and only at 48 h, but with the signal weaker in the radicle cortex cells (Fig. 3b, Supplementary Tables 3, 4, Extended Data Fig. 2). This observation corresponded with our annotation of cluster 14 from known marker transcripts, which indicated the presence of epidermis, endodermis, atrichoblast and root cap marker transcripts (Fig. 2c). The marker gene validation results for both clusters were also consistent with the changes in detection of clusters over time described above, further illustrating the dynamic nature of cell transcriptomes during germination (Fig. 2c, Extended Data Fig. 1e).

**Fig. 3.**
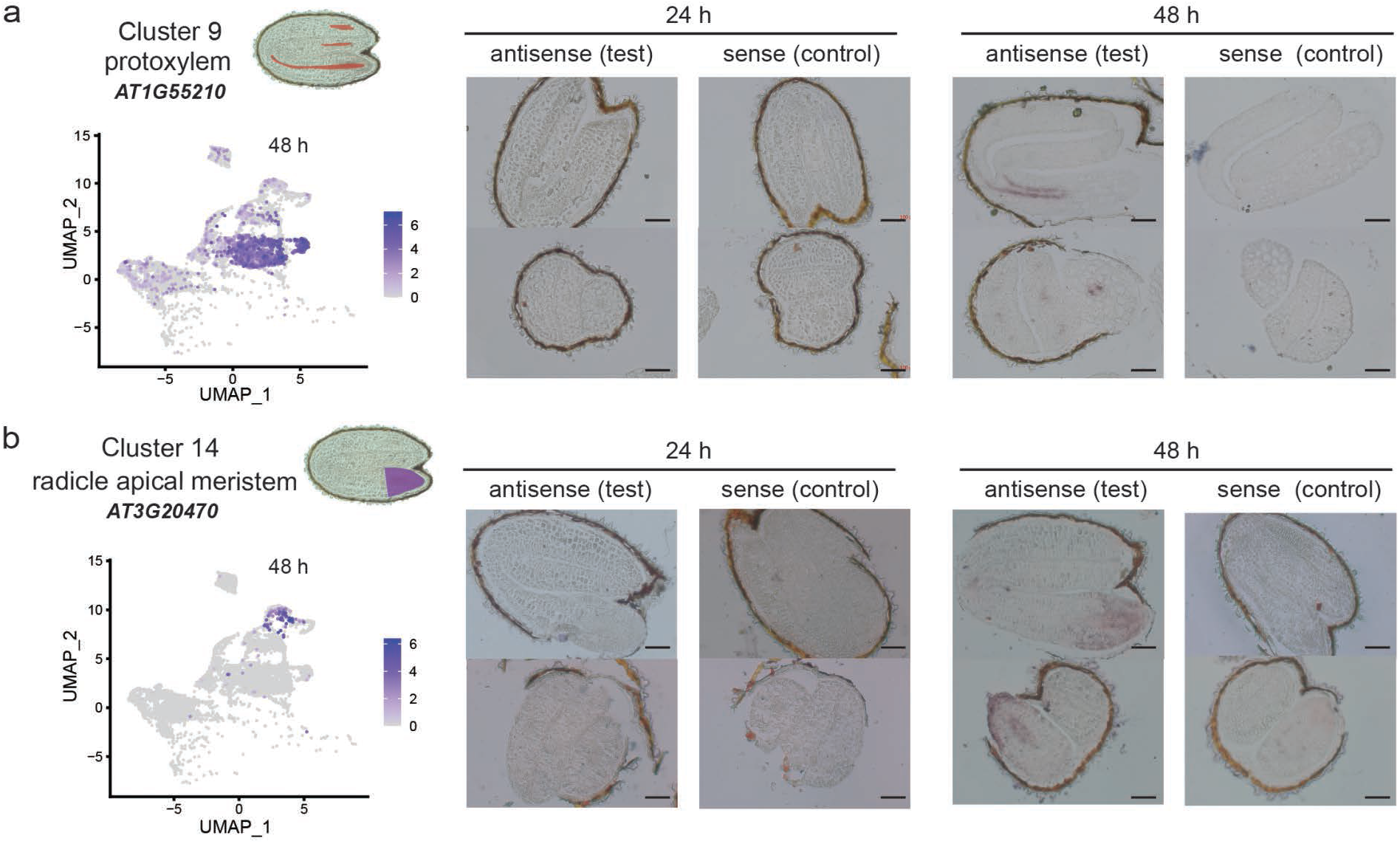
Validation of cell type annotation using RNA *in situ* hybridization. Expression domains of independent marker transcripts of **a,** cluster 9 and **b,** cluster 14 confirmed the physical location of the cells in these clusters. Signals were detected at 48 h, but not at 24 h, confirming also the temporal detection of these clusters in scRNA-seq data. At each timepoint the left panel shows the results of hybridization with antisense probes (i.e. the test), whilst right panels show the results of hybridization with sense probes (negative control). Scale bars indicate 200 μm.

### An initial cell transcriptional state is established early in germination

We sought next to understand how initial cell transcriptional states are established as cells commence activity. The earliest germination time point, 12 h, was dominated by cells of clusters 8 and 11 (cluster 8, 26.74%, and cluster 11, 37.27%, of cells captured at 12 h) (Fig. 4, 1f). Like many other clusters, the presence of clusters 8 and 11 was dynamic across germination, being greatest at 12 h, and with the clusters being almost entirely absent by 48 h (Fig. 4a). We were unable to identify known cell-type marker transcripts from published literature with which clusters 8 and 11 could be annotated. Cluster 8 did express marker transcripts of mature embryos/dry seeds, suggesting that it might be comprised of cells in an early germination state that has not previously been characterized (Fig. 2c) [35]. To test this idea we assessed the similarity of the cluster 8 and 11 transcriptomes with the transcriptomes of whole seeds at earlier germination time points (Fig. 4b). To do so we used a dataset of whole (bulk) seed RNA-seq that included 1, 6 and 12 h germination time points, earlier than in our scRNA-seq, and calculated identity (module) scores between each cell and the bulk seed transcriptomes [13]. Cluster 8 cells identified strongly with transcriptomes of 1 and 6 h bulk seeds, more strongly than all other clusters, whilst cluster 11 did not. This suggests that the biological properties of clusters 8 and 11 are distinct.

**Fig. 4.**
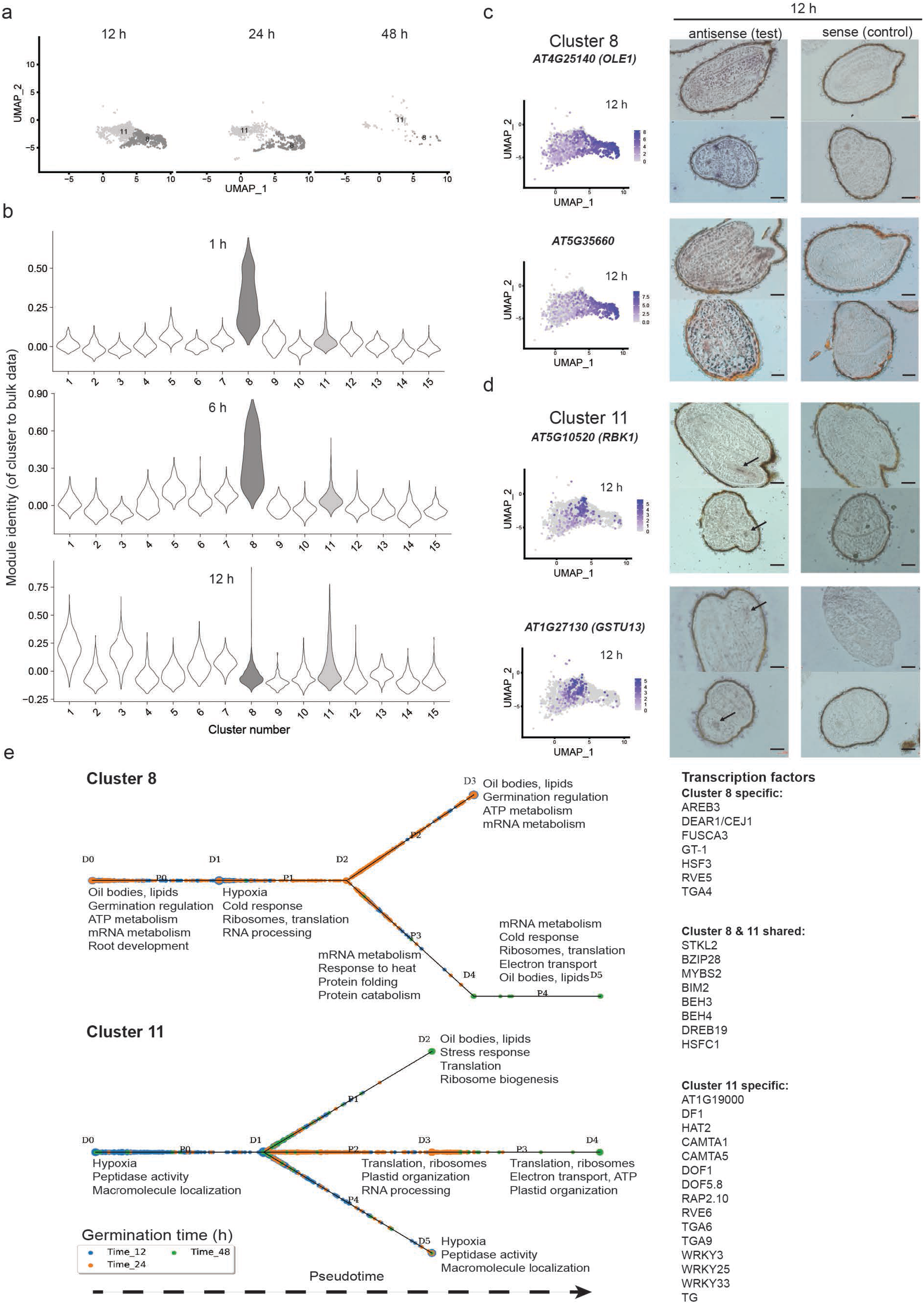
Initial transcriptional states are established early in germination. **a,** UMAP dimensional reduction and visualisation of cluster 8 and 11 cells across three time points, 12 h, 24 h and 48 h. **b,** Comparison of transcriptomes of each cell, grouped by cluster, against whole (bulk) seed transcriptomes from early time points during germination. The transcriptomes of cluster 8 cells are strongly similar to transcriptomes of bulk seeds at 1 and 6 h of germination. **c,d** RNA *in situ* hybridisation to confirm the location of clusters 8 **c** and 11 **d** using two cluster-specific marker transcripts for each. Expression of each marker is shown in an adjacent UMAP dimensional reduction plot; these plots display all cells detected at 12 h, some of which belonging to other clusters. Scale bars = 200 μm. Arrows indicate regions where signals were detected. **e,** Pseudotime models of cell developmental trajectories for clusters 8 and 11, constructed using CSHMM-TF. P indicates paths, D indicates split nodes. Split nodes are the start and end of each path. Major gene ontology terms associated with each stage of models are summarised and predicted regulatory transcription factors listed.

The physical locations of cells in clusters 8 and 11 were determined in order to better understand their biological properties. We examined the localisation of two marker transcripts for each cluster using RNA *in situ* hybridisation. Expression of cluster 8 marker transcripts (*AT4G25140*, *AT5G35660*) was detected throughout the whole embryo (cotyledon, hypocotyl, radicle, and provasculature) at 12 h (Fig. 4c, Supplementary Table 4, 5, Extended Data Fig. 3). Contrastingly, both marker transcripts of cluster 11 (*AT5G10520* and *AT1G27130*) were detected only at the provasculature cells (Fig. 4d, Supplementary Table 4, 5, Extended Data Fig. 3). The expression domain of cluster 11 marker transcripts corresponds to a defined region of abscisic acid and giberellic acid signalling that are proposed to regulate the decision to germinate in dormant seeds, an event that precedes the germination events covered by our experiments [4]. Considered together, these data indicate that cluster 8 represents a general cell transcriptional state through which most cells of the embryo pass early in germination. By contrast, cluster 11 likely represents the set of cells where the decision to germinate was made, which appear to have a different transcriptional state during early germination than all other cells.

As clusters 8 and 11 represent two different populations of cells, we expected that active biological functions would differ between their respective cells. We investigated these biological functions by assembling the cluster 8 and 11 cells onto pseudotime trajectories and analysing the functions of the genes expressed in each cluster using an approach called Continuous-State Hidden Markov Models TF (CSHMM-TF) (Fig. 4e, Extended Data Fig. 4, Supplementary Tables 6, 7) [36]. The CSHMM-TF method assigns activation time of TFs based on both their expression and the expression of their target genes. Not all cells become active at the same time during germination, and there is variability in the precise time of germination between genetically identical seeds [5, 37, 38]. Consequently, the cells will be spread across a developmental trajectory of gene expression, with each cell in a slightly different expression state. Assembly of the cells onto a pseudotime trajectory arranges the cells according to their expression states, and thereby developmental progression, enabling more precise examination of how gene expression changes during germination. CSHMM-TF also identifies where gene expression of groups of cells diverge substantially during pseudotime, splitting cells with different expression states onto different paths.

Cells of cluster 8 first expressed genes involved in utilisation of energy resources (ATP, oil bodies, path P0), followed by RNA processing, translation and hypoxia (P1) (Fig. 4e, Extended Data Fig. 4, Supplementary Tables 6, 7). The expression of energy biology functions earliest likely indicates the initiation of metabolism, whose resumption in low oxygen conditions would result in the observed hypoxia response at this phase of germination [39]. Cells then split along two gene expression paths (P2 and P3/4). In both paths, cells expressed genes involved in mRNA metabolism and energy biology, but path P3/4 cells expressed more protein processing and translation functions. The pattern of gene expression differed in cluster 11 cells compared with cluster 8 (Fig. 4e, Extended Data Fig. 4, Supplementary Tables 6, 7). Genes associated with hypoxia were already expressed at the earliest phase of the model, which would be consistent with metabolism in these cells having become active earlier or more rapidly than cells of cluster 8 (P1). Cluster 11 cells then split along three gene expression paths (P1, P2/3 and P4). Expression of functions involved in translation and ribosome biogenesis featured in two cluster 11 paths (P1 and P2/3), and overall more clearly so than in the cluster 8 model, suggesting translational activity may be greater in cluster 11 cells. For both cluster 8 and 11, the pseudotime arrangement was consistent with the actual germination time of cells (Fig. 4e). Overall, these analyses may indicate that cells of cluster 11 have progressed to a more advanced stage of germination than cells of cluster 8.

Gene expression is regulated by the action of transcription factors that form gene regulatory networks. The differing patterns of gene expression between cells of clusters 8 and 11 indicate different gene regulatory networks act within the clusters. We examined these gene regulatory networks by identifying candidate transcription factors active in cells of either cluster (Fig. 4e, Supplementary Tables 6, 7). CSHMM-TF models also predict which transcription factors are active within each gene expression path of the pseudotime trajectories. This is achieved by integrating transcription factor binding data, here provided as experimentally-validated target genes for >500 Arabidopsis transcription factors from genome-wide in vitro protein-DNA binding assays [40]. Twenty-two predicted regulatory transcription factors were unique to one cluster (7 in cluster 8, 15 in cluster 11), whilst 8 were shared between both clusters. Known regulators of brassinosteroid hormone responses were amongst the transcription factors shared between clusters (BIM2, BEH3, BEH4), which is notable because brassinosteroid has an important role in cell division and growth [41]. Two transcription factors specific to cluster 8 may have functions in embryo development, seed maturation and lipid accumulation (AREB3, FUSCA3) [41–43]. Another cluster 8 specific transcription factor (GT-1) may promote light responsive gene expression, consistent with the recent exposure of the seeds to light as a germination trigger [44]. A cluster 11-specific transcription factors has a role in mucilage production (DF1), which occurs early in germination [45]. Other cluster 11-specific transcription factors are involved in calcium signalling (CAMTA1, 5) and auxin-mediated morphogenesis (HAT2), both signalling pathways that influence seed development and germination [46–50]. These analyses indicate that cells of cluster 8 and 11 are executing distinct gene regulatory programs, involving different sets of transcription factors.

### Embryo cells undergo extensive transcriptional reprogramming as germination progresses

The growth properties and development of individual cells within the embryo change as germination progresses [5, 6]. We investigated how underlying dynamic gene expression might contribute to changes in the functional properties of cells during this time. We focused on the hypocotyl cortex cells because these were detected as three distinct clusters (1, 5 , 7) present in different proportions at each timepoint across germination (Fig. 5). Cells of cluster 5 were the most abundant at 12 h, accompanied by a small number of cluster 1 cells (Fig. 5a). Cells of 1 and 7 were most abundant at 24 h, whilst at 48 h very few cells were detected from any of the clusters but cells of cluster 1 were most abundant. The existence of three distinct clusters of hypocotyl cortex cells indicated that there were populations of hypocotyl cortex cells whose transcriptomes differed. It was not possible to identify single marker transcripts clearly specific to individual clusters, suggesting that the differences between clusters were relatively subtle and related to quantitative differences in expression of many genes rather than complete presence/absence of certain transcripts (Extended Data Fig. 5, Supplementary Table 8). However, marker transcripts strongly specific to all three clusters 1, 5, 7 combined were readily identified (Extended Data Fig. 6, Supplementary Table 9). *AT4G16410* was a specific marker transcript of clusters 1, 5 and 7 at 12 h and 24 h time points in our scRNA-seq data. Expression of the transcript was detected by RNA *in situ* hybridisation in lower and middle hypocotyl cortex cells at at 12 h and 24 h, confirming the cell type annotation of clusters 1, 5, and 7 (Fig. 5b, Extended Data Fig. 7).

**Fig. 5.**
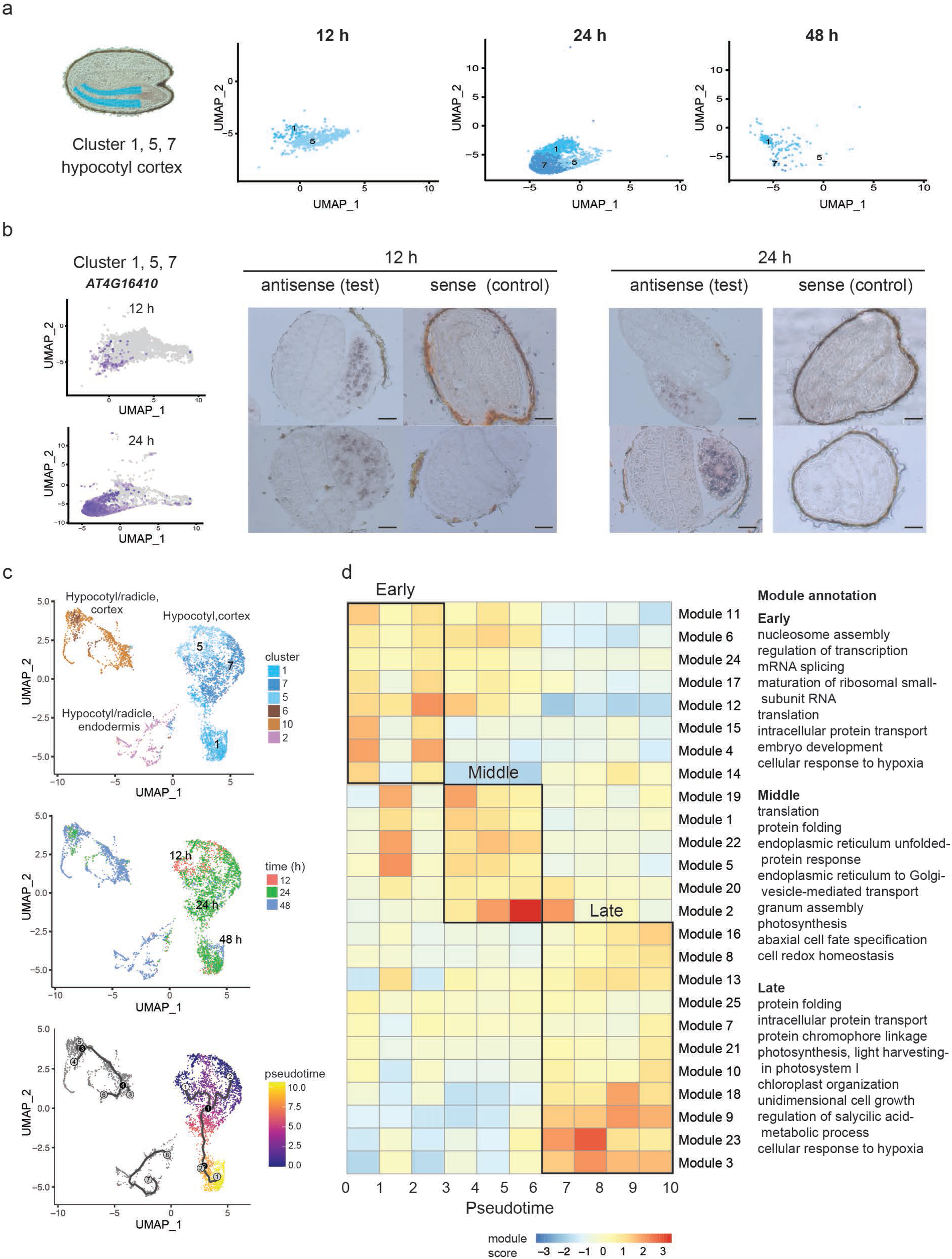
Clusters 1, 5, 7 define a trajectory of hypocotyl cortex cell states. **a,** UMAP dimensional reduction and visualisation of clusters 1, 5, 7 cells across three timepoints, 12 h, 24 h and 48 h. **b,** Confirmation of the location of clusters 1, 5, 7 using RNA *in situ* hybridisation of a marker transcript specific to these clusters at 12 h and 24 h. UMAP dimensional reduction plot of expression of the marker transcript of *AT4G16410* across all cells at each time point. Scale bar = 200 μm. **c,** Cells of clusters 1, 5, 7 form a contiguous group together. They sit upon a temporal trajectory from 12 h to 24 h to 48 h, which corresponds to the transition from cluster 5 cells, through cluster 7 to cluster 1 cells. Reconstruction of pseudotime follows a trajectory that corresponds to the real time of germination. **d,** Co-expressed gene modules across the pseudotime trajectory of cluster 1, 5, 7 cells. Early pseudotime is equivalent to early germination. Module annotations are major representative gene ontology terms associated with modules, assessed using gene ontology term reduction. Complete lists of enriched gene ontology terms are given in Supplementary Table 9.

We conducted a detailed analysis of the transcriptomes of cells within each of the three hypocotyl cortex clusters (1, 5, 7) to understand their similarities and differences. We reclustered and plotted only cells annotated to hypocotyl clusters (1, 2, 5, 6, 7, 10) to remove the influence of the large transcriptional differences from cells of other organs/tissues, thereby allowing us to focus on the smaller differences between hypocotyl cells (Fig. 5c). Cells of the hypocotyl cortex clusters formed a contiguous group even at this focused scale, which transitioned (top to bottom) from cluster 5, through cluster 7 to cluster 1 and indicated that the clusters’ transcriptomes were highly similar (Fig. 5c). There was a corresponding temporal order to the group, transitioning from cells harvested at 12 h (cluster 5), through 24 h (clusters 7 and 1) to 48 h (cluster 1) (Fig. 5a). These observations suggested that clusters 1, 5 and 7 represent hypocotyl cortex cells transitioning through different transcriptional states over time during germination. Reconstruction of a pseudotime trajectory across the clusters supported this proposal, with pseudotime following a similar path to the true time of germination.

The dynamics observed in transcriptomes across hypocotyl cortex cell clusters indicated that the functional properties of these cells change during germination. To examine this we identified modules of genes that are co-expressed across cells, and assessed their functions and timing of expression. There were 25 distinct modules of co-expressed genes across the pseudotime trajectory of clusters 1, 5, and 7 (5d, Supplementary Table 10). These were broadly categorised as early (modules 4, 6, 11, 12, 14, 15, 17, 24), mid (1, 2, 5, 19, 20, 22) and late (3, 7, 8 , 9, 10, 13, 16, 18, 21, 23, 25) on the pseudotime trajectory, which can be considered as equivalent to early, mid and late germination. Genes co-expressed during early germination were enriched for functions related to chromatin accessibility, transcription, RNA splicing, and translation. During mid germination, functions related to translation, protein maturation and photosynthesis were more strongly evident, and in late germination photosynthesis was the dominant function. These analyses indicate that dynamic gene expression drives functional changes in hypocotyl cortex cells across germination.

### Individual gene regulatory programs are active in each cell type of the germinating seed

The many cell types of an embryo each have different roles and contribute at different times to the success of germination [1, 4, 5, 27]. This is achieved by cell types having distinct functional properties, which must be determined by the particular complement of genes these cells express. Consequently, each cell type of the germinating embryo should have a unique and dynamic gene regulatory program. We examined the properties of these gene regulatory programs by reconstructing pseudotime trajectories, using CSHMM-TF, for each cluster identified from our scRNA-seq dataset (Fig. 6, Extended Data Fig. 8, Extended Data Fig. 9, Extended Data Fig. 10, Supplementary Tables 6, 7, 11-13). Gene expression was dynamic and complex across germination in all clusters, with each cluster presenting several modules of co-expressed genes. For example, 13 modules of co-expressed genes were detected in cotyledon mesophyll cells (cluster 13), with a temporal structure of the modules apparent (Fig. 6a). The enriched functional annotations differed between modules, indicating temporal transitions in expressed biological functions. These included translation and chromosome organisation (early pseudotime), hypoxia and photosynthesis (mid), and protein transport/modification (late) (Fig. 6b). Some gene ontology terms were shared between different modules (e.g. modules 3 and 8), reflecting sequential activation of multiple groups of genes that contribute to the same process. The gene regulatory programs of other clusters were different, though they did have similarities. For example, 5 modules of co-expressed genes were detected in protophloem (cluster 15) (Fig. 6c). These modules were associated with translation and temperature/water responses (early pseudo-time), hypoxia and photosynthesis (mid), followed by chromosome organisation (late) (Fig. 6d).

**Fig. 6.**
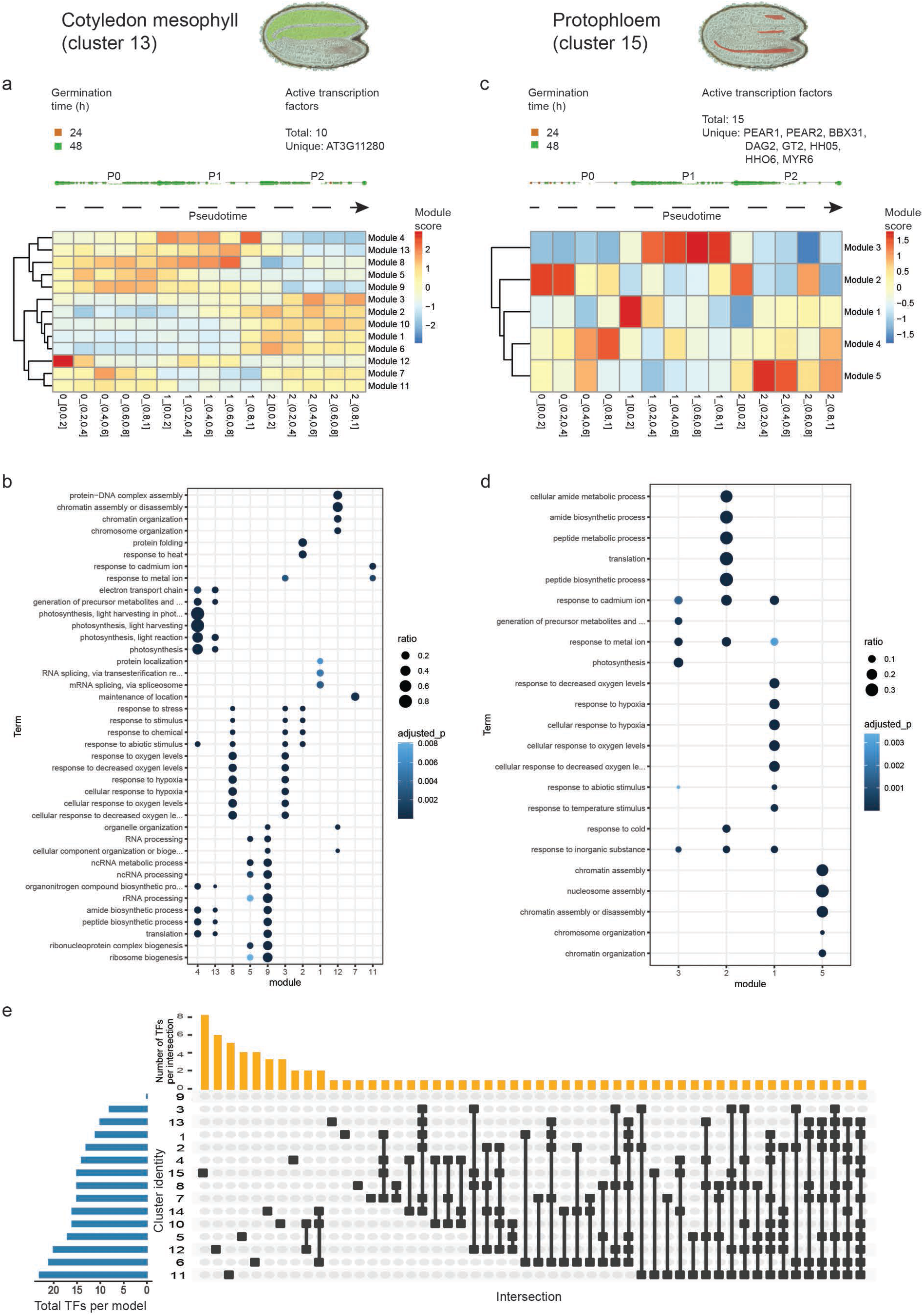
Each cell cluster is defined by a unique gene regulatory program, which reflects dynamic and differing function. **a,** Co-expressed gene modules across the pseudotime trajectory of cluster 13 cotyledon mesophyll cells. **b,** Enriched gene ontology terms of cluster 13. **c,** Co-expressed gene modules across the pseudotime trajectory of cluster 15 protophloem cells. **d,** Enriched gene ontology terms of cluster 15. **e,** Active TFs in every CSHMM-TF model of the 15 clusters identified, comprising a total of 81 unique transcription factors, 39 of which are active in one cell cluster only.

Differing gene regulatory programs would require the activity of different transcription factors. Ten active transcription factors were predicted in the cotyledon mesophyll cell (cluster 13) gene regulatory program, compared with 15 active transcription factors in protophloem cells (cluster 15) (Fig. 6a, b, Supplementary Table 13). Only one of these transcription factors was shared between the two cell types, indicating how the differences in the cell types’ gene regulatory programs may arise. A total of 81 transcription factors were predicted to be active across all cell types (Fig. 6e, Supplementary Table 13). Thirty-nine transcription factors were unique to a single cluster, such as PEAR1 and PEAR2 which were uniquely predicted to regulate the protophloem gene expression modules (cluster 15) and are already known regulators of protophloem development [51]. By contrast, other transcription factors were shared between many cell types, such as BIM2 (a known regulator of brassinosteroid signalling and growth, 8 clusters) and MYBS2 (a known regulator of glucose and abscisic acid signalling, 9 clusters) [41, 52]. Overall, these analyses indicate the different cell types of a germinating embryo express unique and dynamic gene regulatory programs that are likely governed by specific sets of transcription factors.

## Discussion

How cell type specific patterns of gene expression are established and change in individual embryo cells during germination are significant unanswered questions in seed biology. In this study we have comprehensively described gene expression dynamics between 12 and 48 hours of germination in the individual cells of the Arabidopsis embryo. We observe that gene expression is highly dynamic within individual cell types and that cells transition through distinct transcriptional states as germination progresses. Almost all embryo cells pass through the same, single, transcriptional state early in germination, afterwards diverging to their cell-type specific patterns of gene expression. Gene expression dynamics within cell types relate to the functions operating in those cells and are governed by cell-type specific gene regulatory networks. These findings significantly increase our understanding of how gene expression commences during germination. They also provide a general framework within which to study cell-specific gene expression during germination of other Arabidopsis genotypes and plant species where seed strategies differ.

An important insight provided by our study is that the same initial transcriptional state is established in nearly all embryo cells at the start of germination. This was evident from our observations that more than a third of cells at 12 h of germination belonged to a single cluster (8), and that marker transcripts for this cluster were broadly expressed across the embryo when observed using RNA *in situ* hybridization. Similar general transcriptional states through which many cell lineages pass also exist in *Drosophila melanogaster* and humans. Undifferentiated cells in *D. melanogaster* transition through a general transcriptional state in preparation for differentiation [53]. Human cell types can be placed into 5 major categories, each of which is defined by a shared broad transcriptional program [54]. Subsequent highly-specific cell types then arise from these basic programs. Our study indicates that the earliest phase of widespread embryo cell activity during germination is the expression of a shared transcriptional program, from which the many cell-type specific gene expression programs of the embryo emerge. Why cells need to express this shared transcriptional program upon first activity remains to be discovered.

Our study demonstrates that gene expression is highly dynamic and specific within individual cell types during germination. The transcriptomes of each embryo cell type changed substantially as germination progressed, resulting in changes to the molecular pathways and functions expressed by each cell type over time. The hypocotyl cortex cells were an example of this, expressing genes involved in mRNA splicing and transcriptional functions early in germination, progressing to protein maturation and establishment of photosynthesis in mid-germination, and chloroplast organization and cell growth in late germination when the seed-seedling transition occurs. Similar dynamics were observed in every cell type, but in each case the functions expressed and the sequence of changes were specific to the individual cell type. This presumably reflects the unique role of each embryo cell type during germination. Underlying these expression dynamics were cell-type specific gene regulatory networks, defined by groups of transcription factors. Whilst some transcription factors were predicted to be active across multiple cell types, a subset of transcription factors were specific to individual cell types or transcriptional states. This indicates that distinct groups of transcription factors govern the dynamic functional changes of each embryo cell type as germination progresses.

Overall, we illustrate that the cells of the embryo progress through specific transcriptional states as germination progresses. This enables individual cell types to express the genes that define the changing functions of those cell types at the appropriate time, thereby contributing to the successful seed-seedling transition. Seed structures and resources vary remarkably between plant species, requiring different cell types, functions and dynamics. Our study provides a framework for analysis of functional variation in seeds between species and for investigation of how different species establish cell transcriptional states in the embryo.

## Methods

### Plant material and growth conditions

Col-0 and Cvi-0 seeds were sown on MS media plates (containing 3% sucrose). Seeds were sterilised and stratified for 48 h of cold (4°) dark stratification before being transferred to continuous light (at 22°), then collected after 12 h, 24 h and 48 h in the light. Two biological replicates were collected and analysed.

### Dissociation of *Arabidopsis thaliana* embryos into single cells

Seeds were sandwiched between two glass slides and embryos were released mechanically from seed coats by pressing the slides together. Ruptured seed coats and embryos were collected into microcentrifuge tubes and separated from each other using a Percoll gradient. In brief, the samples were resuspended in MC buffer (10 mM potassium phospate pH7.0, 50 mM NaCl, 0.1 M sucrose) and loaded onto a 1 ml 60% Percoll cushion. The tubes were then centrifuged at 800 g for 5 min and the embryo fraction (at the bottom of the tubes) was collected and re-suspended in 0.6 ml MC buffer. A second Percoll gradient was repeated to obtain pellets of embryos without any seed coats. Enriched embryos were cut with razor blades and digested in protoplasting buffer (2% w/v Cellulase, 3% w/v Macerozyme, 0.4 M D-mannitol, 20 mM MES, 20 mM KCl in milli-Q water with the pH adjusted to 5.7 with 1 M Tris/HCL pH7.5, 0.1% w/v BSA, 10 mM CaCl2, and 5 mM β-mercaptoethanol). After 3 hour of digestion, protoplasts were filtered through a 70 μm cell strainer, followed by a 40 μm cell strainer to remove debris, and centrifuged at 500 g for 5 min. The supernatant was removed and the pellet was washed with 2 ml protoplasting buffer without enzymes or mercaptoethanol and centrifuged at 500 g for 5 min. The pellet was resuspended in 50 μl protoplasting buffer without enzymes and mercaptoethanol. Protoplasts were counted using hemocytometer and adjusted to final concentration of 800–1200 protoplasts per μl.

### Bulk RNA-seq library preparation

Col-0 seeds were sown and collected at the 12 h, 24 h and 48 h time points as above, in three independent replicates (batches) of the experiment. Embryos were released from seed coats and enriched by doing Percoll gradient. For bulk RNA-seq, embryos either collected without protoplasting (np) or with protoplasting (p). RNA were extraction using Qiagen RNeasy Plant Mini Kit. RNA quality and integrity were determined using Qubit fluorometer. Libraries were prepared using TruSeq Stranded mRNA Library Prep kit and sequenced by Illumina sequencer Nextseq500 using 75 bp single end kits.

### Bulk RNA-seq analysis

Raw Illumina reads were trimmed for quality and adapter sequences with Trimgalore v0.4.5. Trimmed fastqs were quality checked with FastQC [55], and aligned to the *Arabidopsis thaliana* Col-0 TAIR10 assembly with hisat2 v2.1.0 [56]. Exonic counts aggregated by genes were calculated with FeatureCounts [57] using the Araport11 annotation [58]. Differential expression analysis between the dissociated and non-dissociated embryos was performed in R 3.5.1 [59] with the edgeR package [60, 61]. The design matrix was defined as `model.matrix(~ time + protoplast)`, and glms for each gene were fit with `glmFit`. Genes differentially expressed by the dissociation treatment were identified by performing a likelihood ratio test on the `protoplast` factor with `glmLRT`. We imposed a 1% FDR and a minimum absolute log2-fold change of 1.5 to call genes as significantly induced or repressed by the dissociation.

### Single-cell RNA-seq library preparation

6,000 protoplasts per time point and replicate were loaded onto a Chromium Single Cell instrument (10x Genomics, Millennium Science Australia Pty Ltd, Australia) to generate single-cell GEMs. Single-cell RNA-seq libraries were generated with the Chromium Single Cell 3’ Library and Gel Bead Kit v2 (10x Genomics, Millennium Science Australia Pty Ltd, Australia) according to user guide (Chromium Single Cell 3’ Reagent Kits v2). The resulting libraries were checked on an Agilent 2100 Bioanalyzer, and quantified with the KAPA Library Quantification Kit for Illumina Platforms (Millennium Science Australia Pty Ltd, Australia). The libraries were sequenced on an Illumina Nextseq500 using two 150 bp paired-end kits with 15% PhiX. The raw scRNA-seq dataset was comprised of 26 bp Read1, 116 bp Read2 and 8 bp i7 index reads.

### Single-cell RNA-seq analysis

CellRanger count (v1.3.0) was used to generate raw UMI-count matrices for each sample separately, mapping to the TAIR10 *Arabidopsis thaliana* genome masked for Cvi-0 SNPs with STAR options `--alignIntronMin=10 --alignIntronMax=5000 --scoreDelOpen=−1 --scoreDelBase=−1 --scoreInsOpen=−1 --scoreInsBase=−1` and using the Araport11 AtRTD2 GTF file.

Single cells for the first replicate, containing Cvi-0 and Col-0 cells, were genotyped by first counting the UMIs containing Col-0 or Cvi-0 SNPs for each cell barcode, followed by density-based clustering with DB-SCAN. These steps are included in the sctools package (https://github.com/timoast/sctools). The clustering parameters were optimised for each sample: *ϵ_background_* = 0.5 and *ϵ_margin_* = 0.3 for 12 h rep1, *ϵ_background_* = 0.4 and *ϵ_margin_* = 0.3 for 24 h rep1, *ϵ_background_* = 0.4 and *ϵ_margin_* = 0.15 for 48 h rep1. Cells genotyped as Col-0 were retained for further analysis.

We applied *emptyDrops* [62] from the *DropletUtils* package (v1.6.1) following the guide to distinguish real cells. Further quality control was performed using *scater* (v1.14.6) [63] to remove cells with 1) more than three median absolute deviations (MADs) of the log10 read counts below the median values; 2) more than three MADs of the log10 genes detected below the median. Then, *calculateAverage* was used to remove low-abundance genes with an average count below 0. Genes induced during protoplast isolation were removed before applying the normalization method *calculateSumFactors* with pool-based size factors used from *scran* [64]. Highly variable genes (HVGs) were selected by *modelGeneVar* and *getTopHVGs* with biological variance threshold set as 0. FastMNN [65] was then performed using HVGs to integrate data sets from each sample. MNN dimension reductions were applied to construct a shared nearest neighbour graph with the function provided in *scran*, and the Louvain algorithm from *igraph* [66] was followed to group cells into clusters. MNN dimension reductions were also applied to generate a two-dimensional UMAP for visualization.

### Cluster and cell type annotation

To identify cluster-enriched genes, genes upregulated in each cluster were calculated using *FindMarkers* from *Seurat* (4.0.5) with the P-value < 0.01 [67] (Supp Table 2). The cell type of each clusters was manually annotated and assigned using known cell-type marker genes from the literature (Supp Table 3). Well-characterised endosperm and seed coat marker genes were included to show exclusion of these two tissues and enrichment of embryos in current data.

### Comparison to bulk RNA-seq data

We also compared the scRNA-seq data to the published time-series sequencing bulk RNA-seq profiles of seed germination [13]. The samples used for bulk RNA-seq were collected after 48h of dark stratification, followed by 1 h, 6 h, 12 h, and 24 h of exposure to continuous light. Differential expression analysis between samples from specific time point to others was performed with the *limma* package [68]. With design matrix defined as `model.matrix(~ 0 + time)`, precision weights are calculated by *voom* [69], and used in `eBayes` for statistical testing. The contrasts were made between data from every time point to the average of data from other time points. Genes that are differentially expressed with the log2-fold change above 1.5 and FDR < 5% are retained as DE genes. We then filtered out genes that are regarded as DE genes at more than one time point. Then the lists of unique DE genes of individual time points are used in the scRNA-seq data to calculate their average expression in each of the cells with the function `AddModuleScore` [70].

### RNA *in situ* hybridisation

Seeds were harvested and fixed in ice-cold Farmer’s fixative (3:1 ethanol:acetic acid). The samples were placed in the cold room (4°) overnight. The following day, the fixed tissues were dehydrated using the Leica Semi-Enclosed Benchtop Tissue Processor TP1020 (Leica Biosystems, Mount Waverley, Australia) at room temperature in a graded series of ethanol (1 h each 75%, 85%, 100%, 100%, and 100% v/v). The tissues were then transferred to a graduated ethanol:xylene series (1 h 20 mins each 75%:25%, 50%:50%, 25%:75% v/v), finished with a xylene series (1 h each 100% and 100% v/v). Tissue was then added to molten Surgipath Paraplast® Paraffin (Leica Biosystems) for 2 times for 2 h each at 65°. Paraplast blocks were then prepared with dozens of seeds in each block using the Leica Heated Paraffin Embedding Module EG1150 H with the added Leica Cold Plate for Modular Tissue Embedding System EG1150 C (Leica Biosystems) with vacuum infiltration. Embedded tissues were cut at eight-micrometer sections and *in situ* hybridization was carried out according to a modified protocol from Jackson (1991): 50°hybridization temperature and 0.2x SSC washes [71]. Transcripts of interest were amplified using designed primers (Supplementary Table 4) and cloned into pGEMT-Easy vector (Promega). Digoxigenin-labelled antisense and sense RNA probes were transcribed from T7 or SP6 promoter of pGEMT-Easy vector (Promega) according to manufacturer’s instructions. All hybridization results were observed and photographed using a Zeiss Axio Observer A1 microscope (Carl Zeiss AG).

### Trajectory inference analysis

Monocle 3 (1.0.0) [72] was used to construct single-cell trajectories. Cells from annotated hypocotyl clusters were extracted and re-processed (including normalization and batch effect correction [65], dimensionality reduction, and clustering) with Monocle 3. This resulted in three distinct partitions, and we learned the trajectory for each of the partitions. We selected the beginning nodes of the trajectory where more adjacent cells come from 12 h. Modules of co-regulated genes were then calculated using Louvain community analysis based on genes with the function *find_gene_modules*. Aggregated expression of all genes in each module across cells along pseudotime was plotted in a heatmap. After grouping modules based on their expression pattern according to the pseudotime stage, we assessed enriched gene ontology terms using all genes from each stage. Gene Ontology analysis was performed using the topGo package [73].

### Inferring TF activation order with *CSHMM-TF*

*CSHMM-TF* [74] was used to analysis of time series single-cell expression data with information about transcription factors (TFs) provided (TF binding data from [40]). Cells from each cluster were extracted separately and their raw count matrix and information about their collected time were used as input. *find_gene_modules* was used to find gene modules across individual clusters. Aggregated expression was then calculated based on assigned time blocks of activation along the learned path by *CSHMM-TF*. GO analysis was performed in each module of genes separately, and the results are shown in dot plots with size denoted by the ratio of provided genes by all genes in a specific term and color denoted by adjusted enrichment P-values.

### GO analysis

Gene Ontology analysis was performed using the topGO package [73]. Whole gene sets without protoplasting genes were used as the background. Adjusted P values of genes enrichment were obtained by multiple P values generated from topGO with the number of tests run in each enrichment analysis.

## Data availability

Raw and processed bulk and single-cell RNA-seq data are available at EBI Annotare under accessions E-MTAB-12521 and E-MTAB-12532

## Acknowledgements

Work in M.G.L.’s lab is funded by the Australian Research Council (ARC) Discovery Program grant DP220102840. Q.G. is funded by an NHMRC Investigator Grant (GNT2007996). J.W. and L.C.L. were funded by an ARC Discovery Program grant DP210103258. This study was supported by the La Trobe Genomics Platform. This work was also supported by the following grants to R.L.: National Health and Medical Research Council Investigator Grant GNT1178460, and ARC grants DP210104058, LE170100225, and CE140100008.

## Authors’ contributions

L.C.L, J.W., Q.G. and M.G.L. conceived and planned the project. R.L., T.S. and M.O. provided novel methods and reagents. L.C.L., M.O., M.P-L., S.N., M.T-O., O.B. and U.V.T.H. contributed to sample preparation, protoplast isolation and RNA *in situ* hybridization. L.C.L., A.H. and T.S. carried out library preparation. Y.Y., M.E.R. and Q.G. processed and analysed the data. L.C.L., Y.Y., Q.G., M.G.L. and G.W.B interpreted results and drafted the manuscript. R.L., M.E.R. and J.W. provided critical feedback on data interpretation. All authors reviewed and contributed to the final manuscript.

## Supplementary information

**Supplementary Table 1.** Genes differentially expressed in response to protoplast isolation and which of these are shared with a prior study by Birnbaum and colleagues.

**Supplementary Table 2.** New marker genes per cluster, defined in this study.

**Supplementary Table 3.** Literature curated marker genes used to annotate clusters to cell types.

**Supplementary Table 4.** Transcripts selected for RNA *in situ* hybridization and primers sequences for cloning.

**Supplementary Table 5.** Cluster 8 and 11 specific marker genes.

**Supplementary Table 6.** CSHMM-TF model, active transcription factors and gene ontology analysis for cluster 8.

**Supplementary Table 7.** CSHMM-TF model, active transcription factors and gene ontology analysis for cluster 11.

**Supplementary Table 8.** Marker genes calculated individually for clusters 1, 5, 7 v all other clusters.

**Supplementary Table 9.** Marker genes calculated for clusters 1,5,7 combined v. all other clusters.

**Supplementary Table 10.** Lists of genes associated with each expression module from a pseudotime model integrating the hypocotyl cortex cell clusters 1, 5 and 7. The top 10 gene ontology (ranked by smallest p-value) terms enriched amongst genes of each module are also given.

**Supplementary Table 11.** CSHMM-TF models for all clusters - ordering of cells and gene module membership.

**Supplementary Table 12.** Gene ontology for CSHMM-TF models for all clusters.

**Supplementary Table 13.** Active TFs for CSHMM-TF models for all clusters, plus a list of TFs unique to a single model/cluster.

---

**Extended Data Fig. 1.**
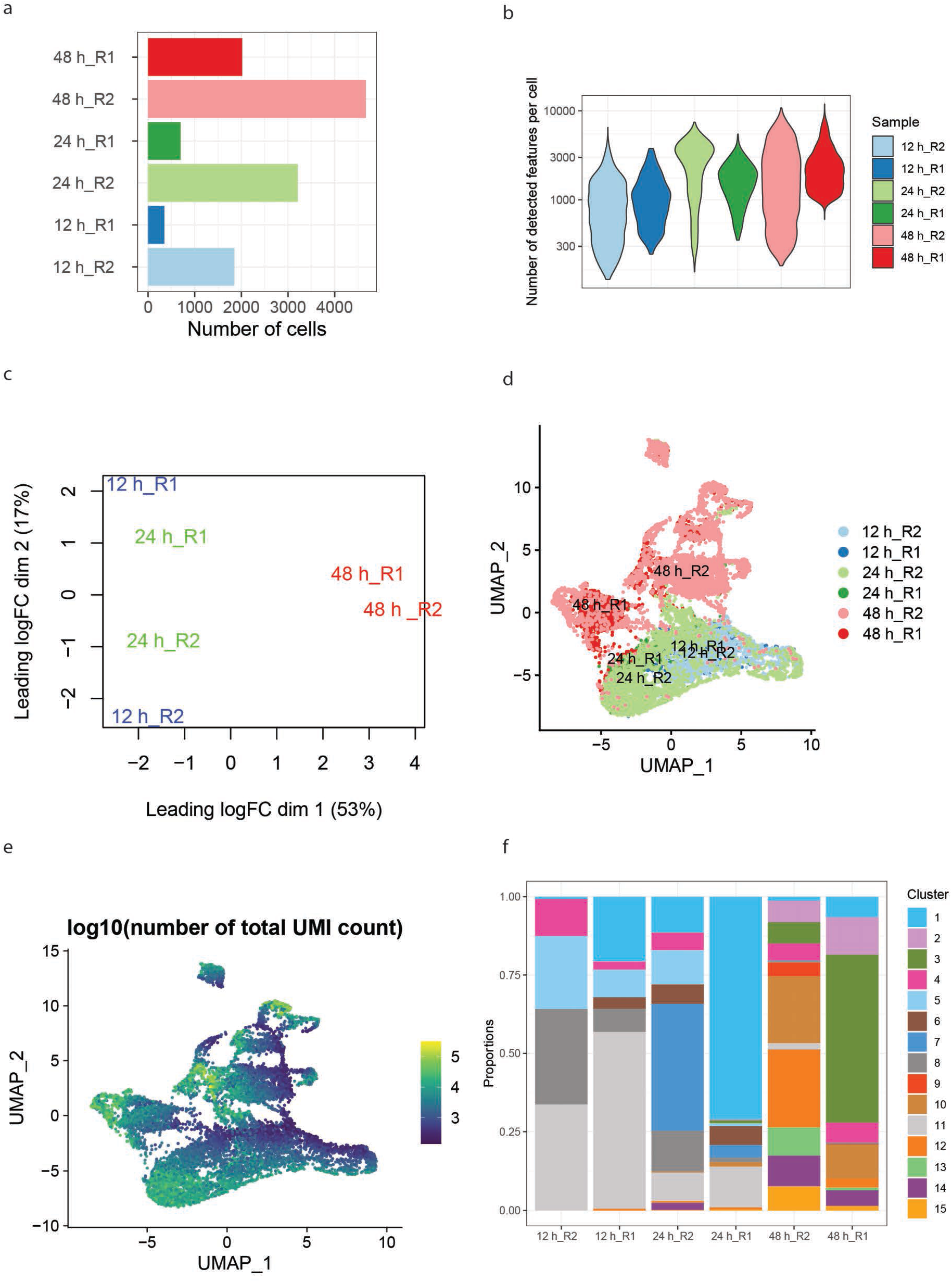
Quality assessment and integration of scRNA-seq data. **a,** Numbers of cells captured for each time point (12, 24, 48 h) and biological replicate (R1, R2). **b,** Distribution of number of detected genes per cell. **c,** MDS plot of pseudo-bulks for each scRNA-seq sample. **d,** UMAP plot depicting relative similarity of all cells post batch correction and data integration. **e,** Total Unique Molecular Identifier (UMI) count per cell for all cells. **f,** Proportional distribution of cells between clusters in each sample.

**Extended Data Fig. 2.**
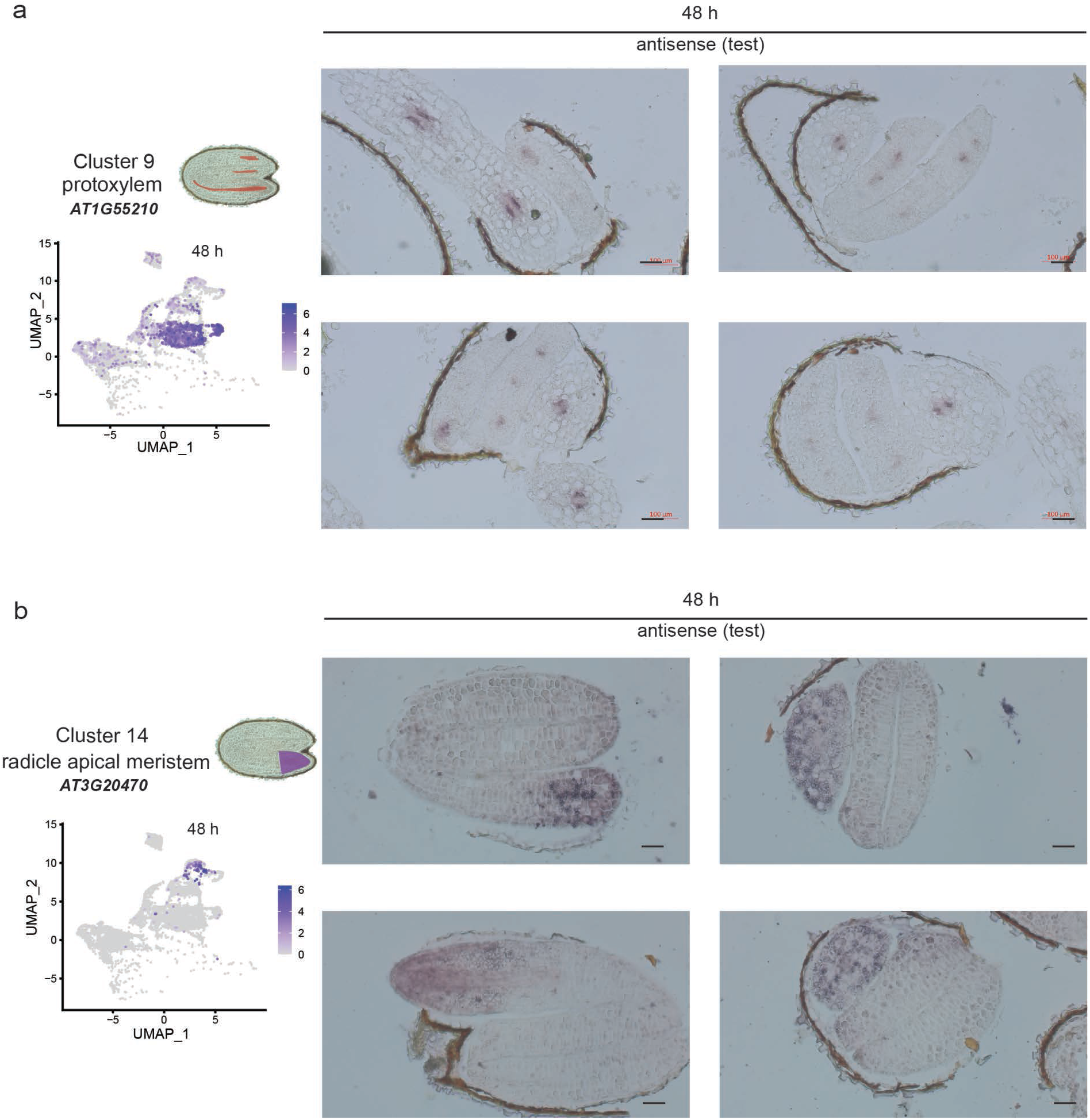
RNA *in situ* hybridization of marker transcripts specific to cluster 9 and cluster 14.

**Extended Data Fig. 3.**
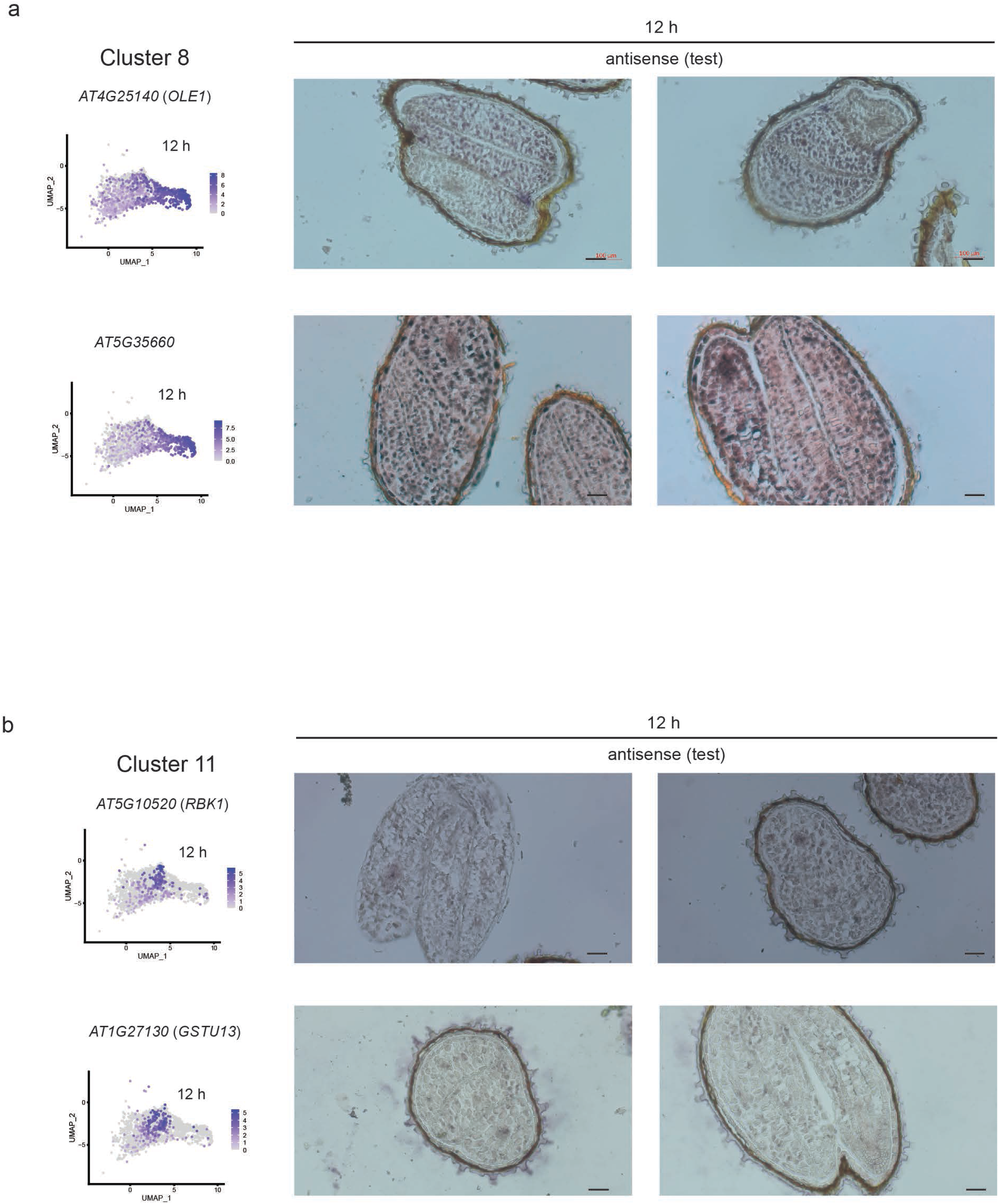
RNA *in situ* hybridization of marker transcripts specific to cluster 8 and cluster 11.

**Extended Data Fig. 4.**
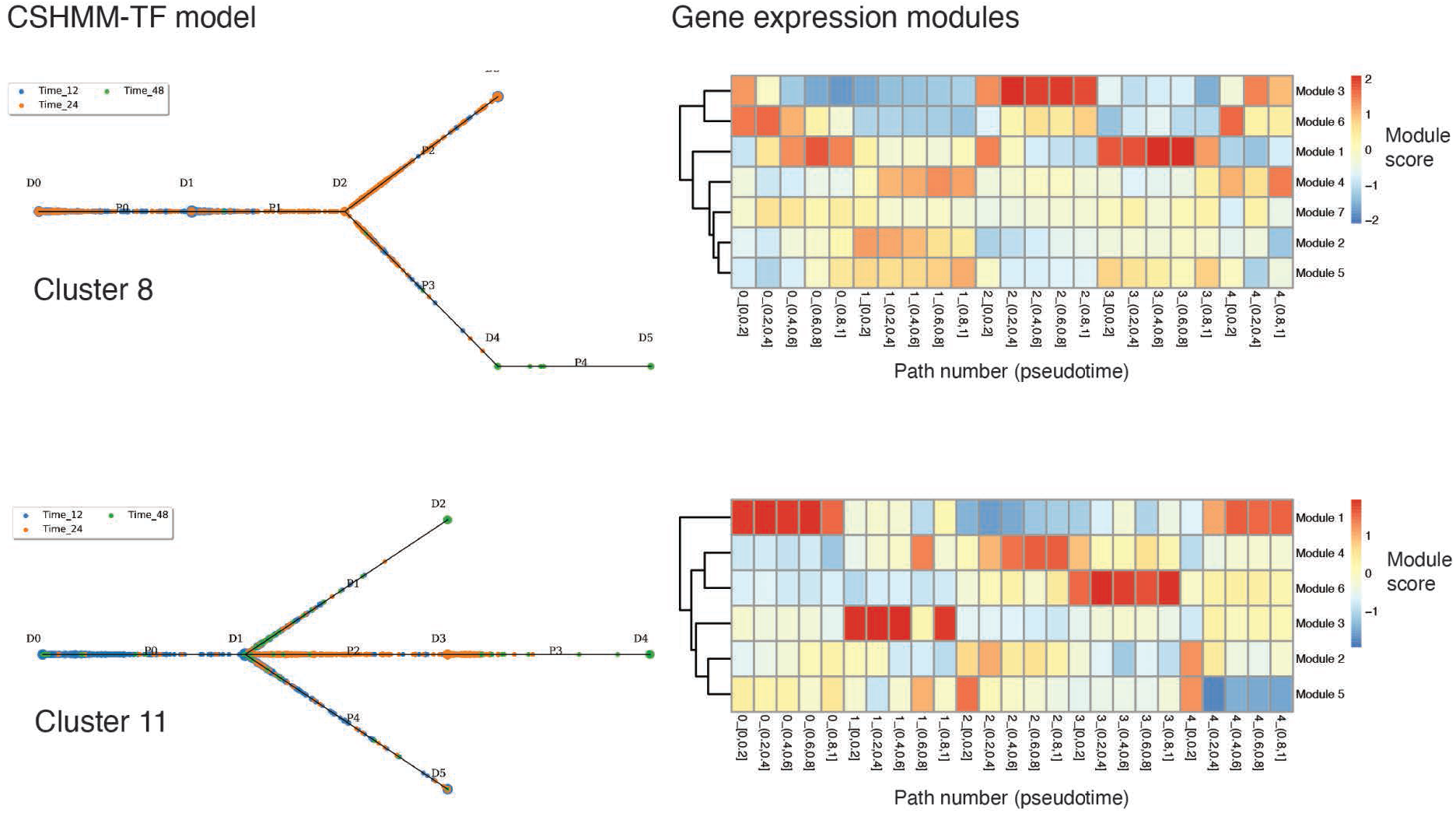
CSHMM-TF model gene expression modules for clusters 8 and 11. Left, CSHMM-TF models. P indicates paths, D indicates split nodes. Split nodes are the start and end of each path. Right, heatmap depicting modules of gene co-expression identified within the CSHMM-TF models. At bottom of co-expression heatmap x-axis labels indicate path number and, in brackets, relative position in pseudotime within the path. Colour scale indicates the module score, which is essentially the average log-fold change of all genes within a module compared to the background control. Functions of genes highly co-expressed within modules were subsequently analysed using gene ontology, to understand what the gene functions characteristic of each module.

**Extended Data Fig. 5.**
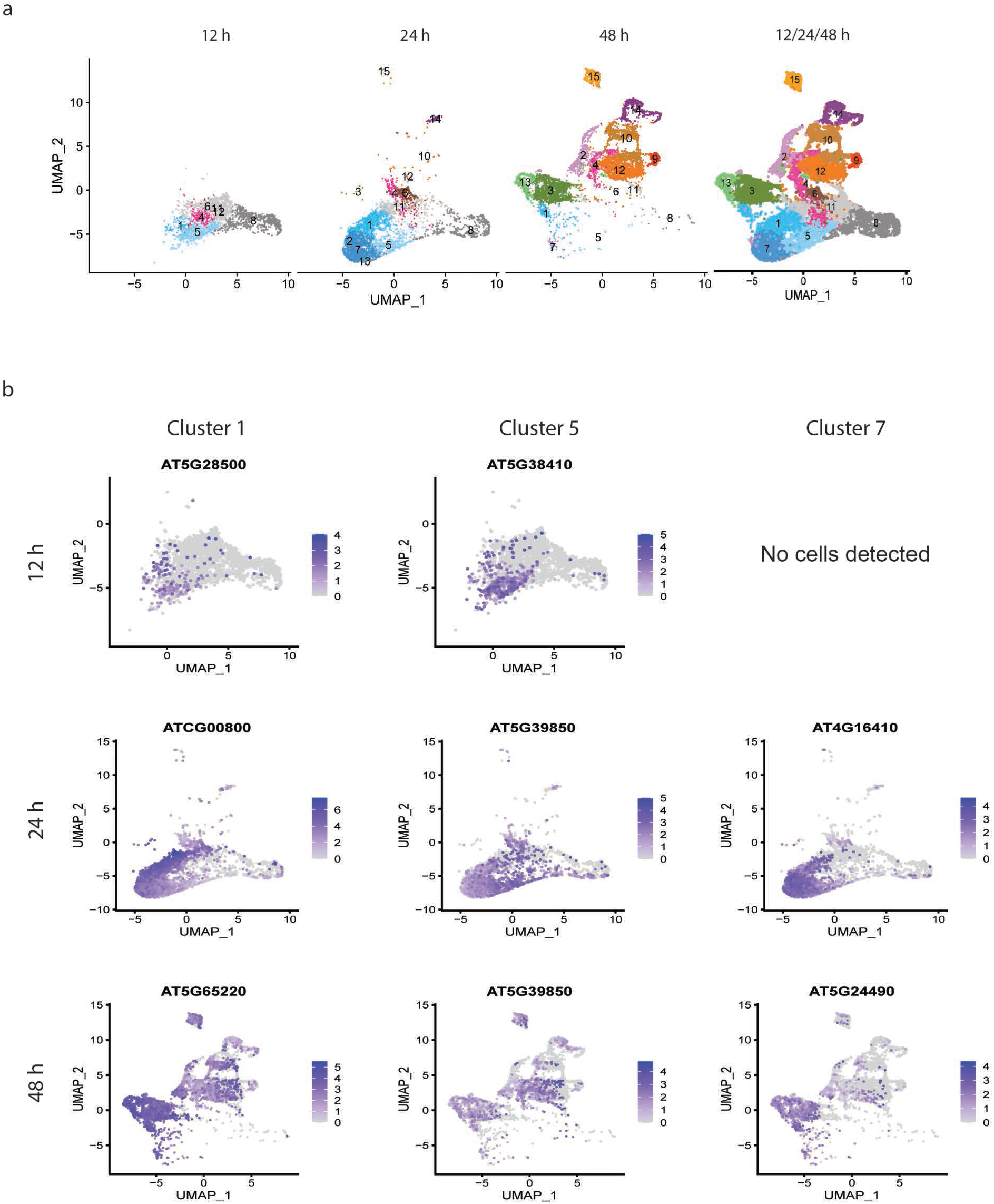
Identification of marker transcripts for clusters 1, 5 and 7. **a,** UMAP dimensional reduction and visualisation of cells at three individual time points, 12 h, 24 h, 48 h, and all time points together. **b,** Most highly specific marker transcripts for each of clusters 1, 5 and 7 individually. The plots illustrate that marker transcripts highly specific to each of these clusters individually could not be identified, likely due to high similarity in transcriptomes between the three clusters. These most highly individual cluster-specific marker transcripts were still expressed in clusters other than 1, 5 and 7.

**Extended Data Fig. 6.**
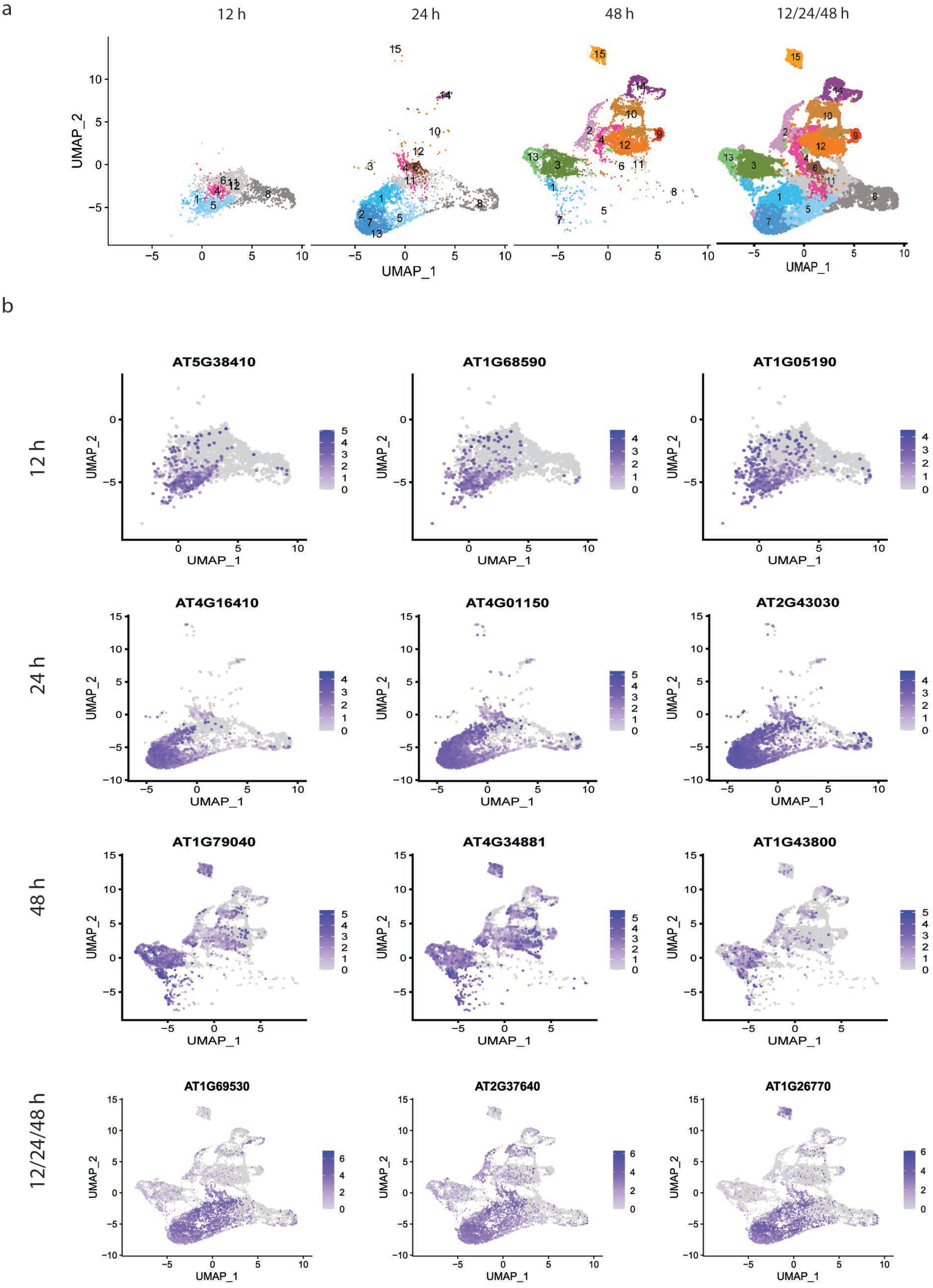
Identification of combined marker transcripts for clusters 1, 5 and 7. **a,** UMAP dimensional reduction and visualisation of cells at three individual time points, 12 h, 24 h, 48 h, and at all time points together. **b,** Most highly specific marker transcripts for clusters 1, 5, 7 combined. Marker transcripts identified were more highly specific to the clusters when these clusters were analysed as a group.

**Extended Data Fig. 7.**
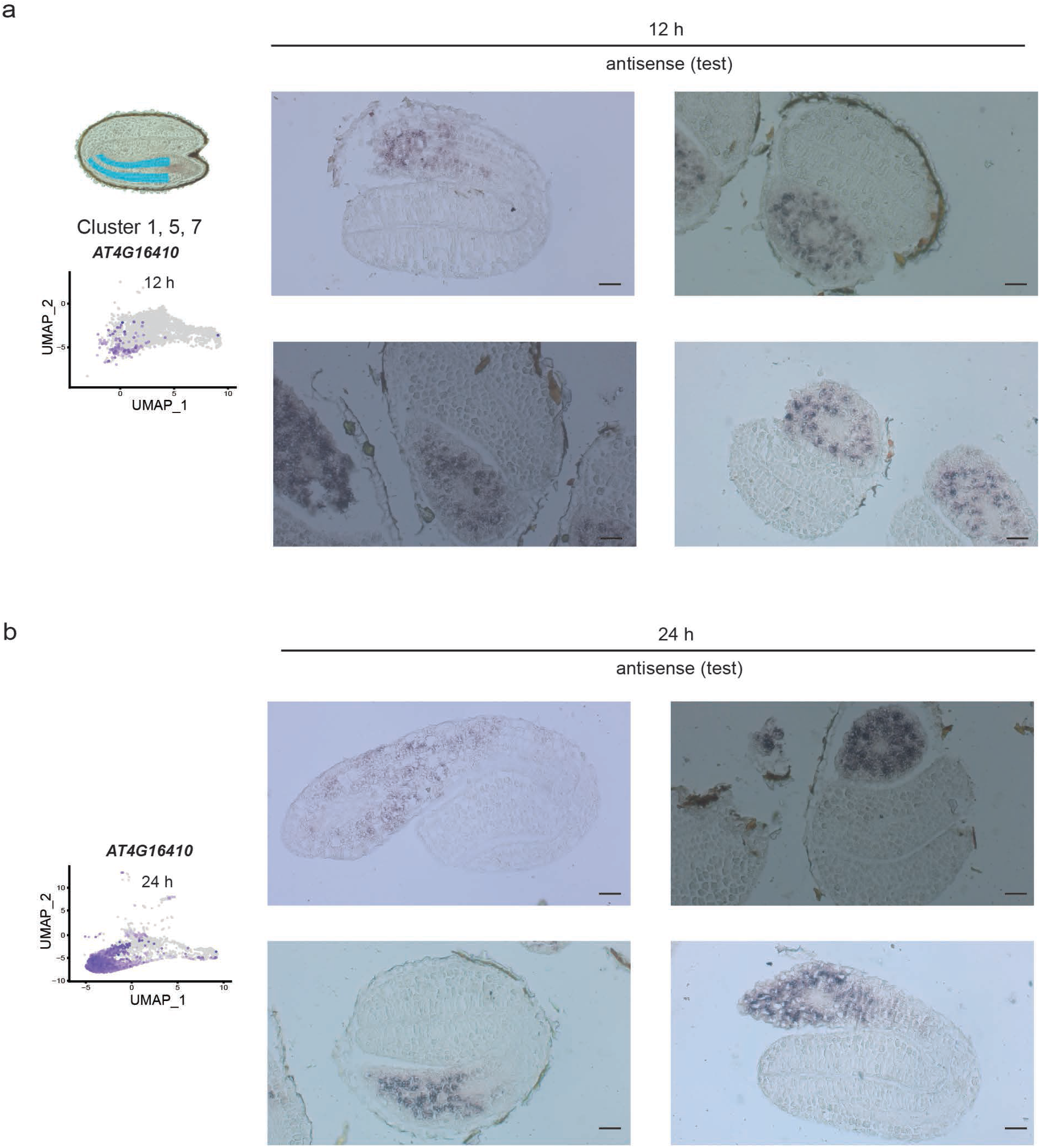
RNA *in situ* hybridization of marker transcripts specific to clusters 1, 5, 7 combined.

**Extended Data Fig. 8.**
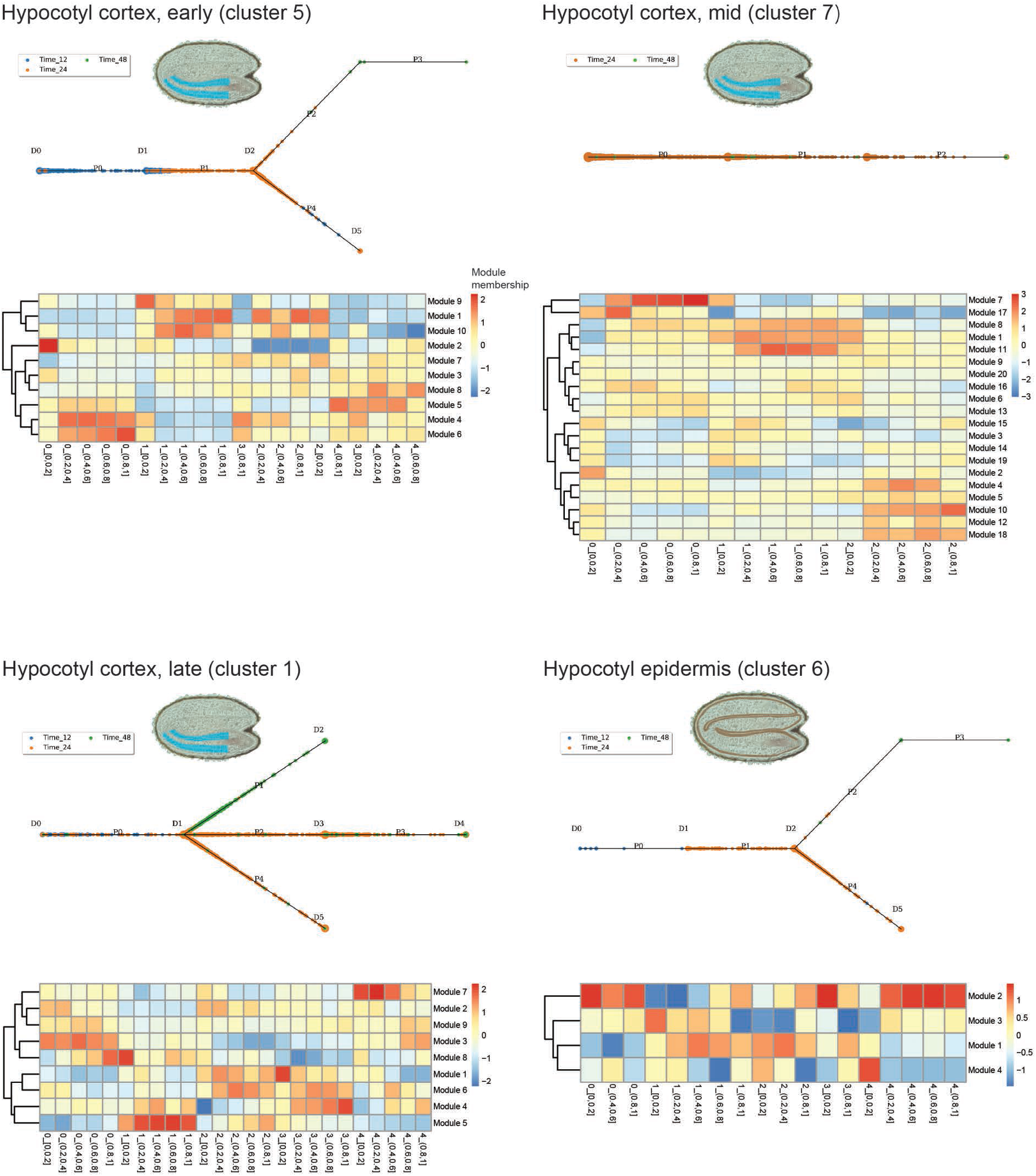
CSHMM-TF models for clusters 5 (hypocotyl cortex, early), 7 (hypocotyl cortex, mid), 1 (hypocotyl cortex, late) and 6 (hypocotyl epidermis).

**Extended Data Fig. 9.**
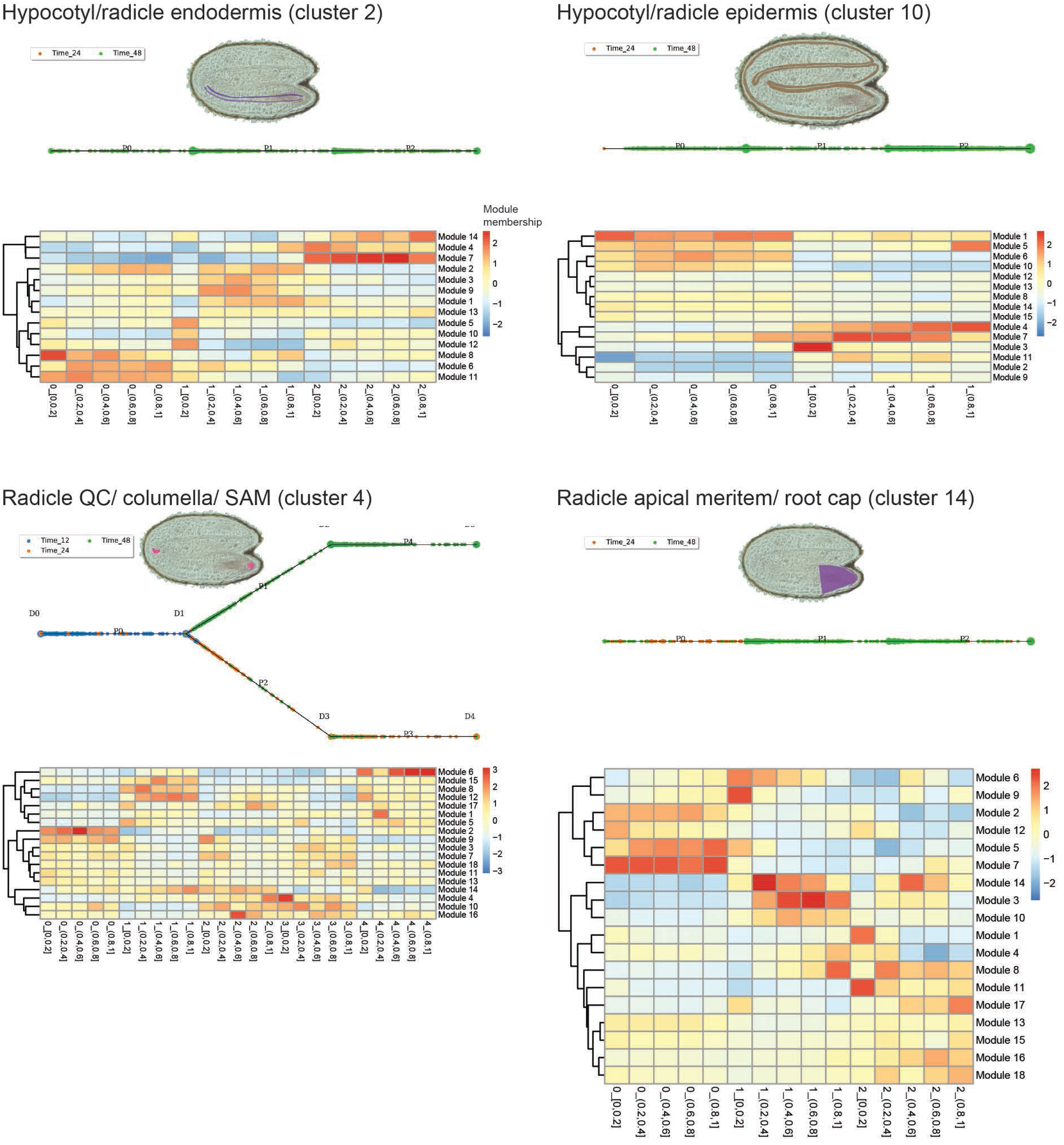
CSHMM-TF models for clusters 2 (hypocotyl/radicle endodermis), 4 (radicle quiescent centre - QC, shoot apical meristem - SAM, columella), 10 (hypocotyl/radicle epidermis, and 14 (radicle apical meristem region).

**Extended Data Fig. 10.**
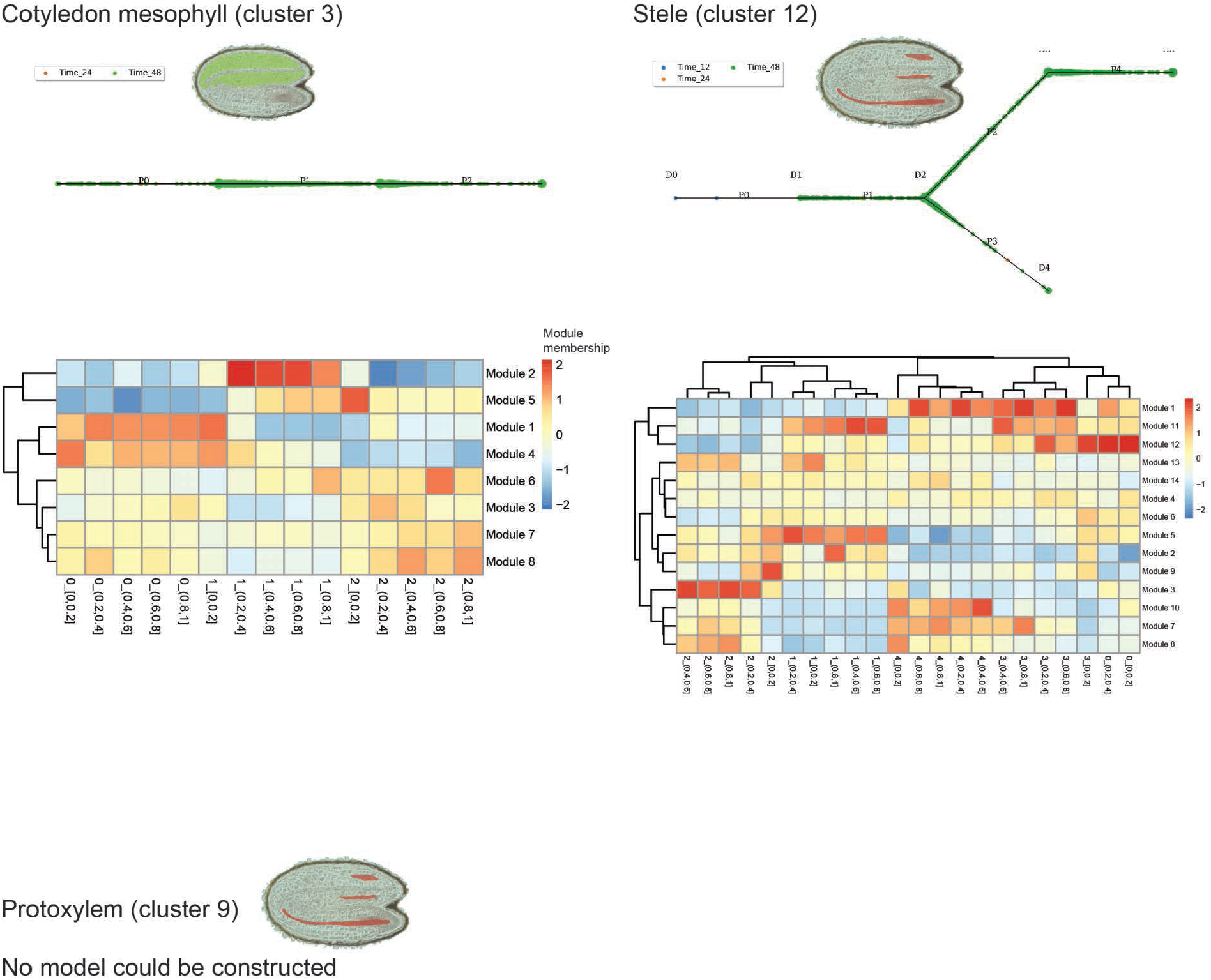
CSHMM-TF models for clusters 3 (cotyledon mesophyll), (12) stele, and protoxylem (9).

